# Constructing Gene Regulatory Network using Chatterjee’s Rank Correlation with Single-cell Transcriptomic Data

**DOI:** 10.1101/2025.09.17.676530

**Authors:** Shreyan Gupta, Anamitra Chaudhuri, Vishnuvasan Raghuraman, Yang Ni, James J. Cai

## Abstract

Constructing gene regulatory networks (GRNs) from single-cell RNA sequencing (scRNA-seq) data is critical for understanding cellular systems, yet existing methods rely either on strong modeling assumptions or on computationally intensive algorithms. We introduce a multiple testing framework for GRN construction based on Chatterjee’s rank correlation coefficient, a nonparametric measure of dependence that captures non-linear and non-monotone relationships. The framework overcomes several limitations of traditional methods by offering a transparent, scalable, and computationally efficient alternative to recent black-box machine learning models. Because cells in scRNA-seq data are not independent observations, we develop a data-driven algorithm for estimating robust testing cutoffs. We further exploit the asymmetric nature of Chatterjee’s correlation to propose a new test for asymmetric predictive associations, yielding biologically meaningful and directionally informed GRNs. In simulations, our method matches or outperforms state-of-the-art approaches in recovering true gene-gene dependencies and asymmetric associations, particularly for complex, non-linear dependencies. Application to real scRNA-seq data shows its utility for dissecting complex regulatory networks.

## 1 Introduction

Deciphering cellular behavior relies on understanding the intricate network of molecular signals that orchestrates cell function. A good example is the response to growth factors, where a signal propagates through a cascade of receptor activation and phosphorylation events, ultimately altering gene expression by activating transcription factors [1, 2]. Such signaling cascades highlight a fundamental principle of cell biology: that regulation is inherently directional, and accurately mapping these directed relationships remains a key challenge [3]. Reverse engineering of gene regulatory networks (GRNs) from transcriptomic data offers a powerful computational approach for understanding gene interactions more broadly. Single-cell RNA sequencing (scRNA-seq) has further enabled GRN inference at a resolution beyond what bulk RNA-seq can offer [4], allowing us to dissect the regulatory logic that governs cell fate, differentiation, and disease, and thereby offering deeper insights into biological systems at a cellular level.

Although several existing methods have been proposed to construct GRNs, they suffer from key limitations that restrict their effectiveness and interpretability. Traditional correlation-based methods, such as Pearson’s and Spearman’s correlations, rely on restrictive assumptions, such as linearity or monotonicity, which rarely hold in complex biological systems [5], and, being symmetric, they cannot capture directional dependence between genes. Other nonparametric measures of dependence, such as distance correlation [6] and mutual information [7], relax these distributional assumptions and can capture nonlinear relationships, but remain inherently symmetric and therefore cannot distinguish which gene is more predictive of the other; mutual information is further hampered by the need for density or binning estimation, which becomes unstable and computationally costly in the sparse, high-dimensional setting of scRNA-seq data. More recent machine learning (ML) approaches like GRNBoost2 [8] and PCNet [9], though flexible and capable of capturing asymmetric relationships, are computationally intensive and often function as black boxes, lacking transparency in how regulatory links are inferred [10]. Moreover, nearly all of these methods, whether correlation-based or ML-based, implicitly assume that cellular observations are independent and identically distributed (IID). The IID assumption is routinely violated in scRNA-seq data, where cell-cell signaling, shared differentiation trajectories, and other biological cross-talk induce dependencies among neighboring cells, undermining the theoretical guarantees on which these methods rely.

To address these combined challenges, we propose a new GRN construction method based on Chatterjee’s correlation coefficient [11], a nonparametric measure of dependence that is asymmetric by design. We additionally evaluate the nearest-neighbor extension of this coefficient proposed by Lin and Han [12], which incorporates multiple right nearest neighbors in the rank construction to boost the power of the original statistic against weak dependence, while retaining its asymmetry and distribution-free nature. While Chatterjee’s correlation has found successful applications in diverse fields, including geochemical sciences [13] and the inference of gene dependencies along scRNA-seq-derived differentiation trajectories [14], its potential for detecting asymmetric gene-gene dependence in scRNA-seq, particularly when the IID assumption is violated, remains largely unexplored. Our approach (a) does not require distributional assumptions, (b) captures directional (asymmetric) dependence between genes, unlike distance correlation or mutual information, (c) can accommodate both linear and non-linear dependencies, (d) remains interpretable through rank-based construction, (e) is computationally simple and scalable, and (f) relies only on local information, i.e., it evaluates each gene pair independently, unlike ML models, which require global modeling.

Although Chatterjee’s coefficient has shown growing theoretical promise, current studies rely on the assumption of independent observations, which fails in the context of gene expression captured by scRNA-seq, where dependencies exist among cells due to shared biological origins, differentiation fates, and cellular cross-talk. To address this gap, our second contribution is an algorithm for estimating empirical cutoff values for testing dependency, based on synthetic or curated datasets with known ground truths. These cutoffs allow practical application of the method to real-world datasets, and in simulations we show that our approach is competitive with substantially more complex ML models and outperforms classical and other nonparametric pairwise measures in recovering nonlinear dependencies. Finally, we use the asymmetric nature of Chatterjee’s correlation to introduce a test for asymmetric predictive association, determining whether the expression of one gene is more nearly a function of another gene than vice versa. Such asymmetry is compatible with, but does not establish, a regulatory direction, as it can also arise from non-invertible relationships, unequal noise, latent confounding, or cell-state mixtures; we therefore interpret the resulting edges as putative predictor → response relationships rather than causal regulation. To the best of our knowledge, this directional test has not been addressed in prior literature, enabling us to construct directed GRNs that prioritize candidate regulatory relationships. Empirically, we demonstrate that our method detects asymmetric association with higher sensitivity than GRNBoost2 in simulation and generates interpretable, testable hypotheses when applied to real single-cell datasets.

An alternative approach to constructing GRNs is via directed acyclic graphs (DAGs) [15]. However, DAG learning suffers from identifiability issues and computational complexity due to the super-exponential growth of the DAG space with the number of genes [16, 17]. In contrast, our framework focuses on pairwise relationships: we first identify dependent gene pairs through multiple testing procedures and subsequently test for asymmetric association at the pairwise level. This correlation-based strategy avoids the computational burden of global DAG search while still yielding biologically meaningful insights. Thus, while DAG-based approaches provide a principled framework for global causal discovery [18–22], our method offers a complementary perspective by prioritizing scalability and interpretability through pairwise testing. In this sense, our framework can also be utilized as a screening approach by efficiently filtering for potentially relevant gene pairs through pairwise testing, which can then be used as inputs for DAG-learning approaches, substantially narrowing the search space and thereby alleviating some of the computational and statistical burdens inherent to global DAG learning.

In the following sections, we introduce Chatterjee’s correlation coefficient as a new approach for identifying gene-gene dependencies and asymmetric (directional) dependence. We first present a data-driven method for establishing cutoffs that effectively control per-null false positive rates (FPRs) and false discovery rates (FDRs), which is crucial for a fair comparison of our method against established correlation, regression, and ML-based approaches. We then evaluate Chatterjee’s correlation against a broad spectrum of established methods for GRN construction. To make our method applicable to real-world scenarios, we propose a technique for estimating cutoffs from prior knowledge and demonstrate its transferability across different scRNA-seq datasets with similar data characteristics. Finally, we apply our approach to real single-cell data, constructing directed GRNs whose edges represent putative predictor → response relationships. This work establishes a scalable and interpretable tool for GRN inference that generates directional inferences about the complex regulations that govern cell fate decisions.

## 2 Results

### 2.1 Chatterjee’s Correlation Accurately Infers Gene Dependencies in Complex Simulated Data

The high cellular resolution of gene expression captured by scRNA-seq is particularly advantageous for reverse engineering GRNs based on co-expression patterns on an intra-sample basis. Furthermore, the substantial number of cells yielded by scRNA-seq, as well as diverse patterns of gene-gene interaction, favors the application of non-parametric statistical coefficients, such as Chatterjee’s correlation coefficient, and we investigate its efficiency in the following studies. Specifically, we conduct two separate studies: first, we perform our experiments on in-house simulated data, and second, we apply our methods to simulated scRNA-seq data using SERGIO [23].

#### 2.1.1 First study: Benchmarks on simulated scRNA-seq data

To comprehensively evaluate the performance of all proposed methods under a range of different gene interaction patterns, we simulated scRNA-seq data incorporating both linear and non-linear gene interactions to mimic real datasets (detailed in Section 3.3). Our simulation used a Poisson approximation and four distinct types of genegene interactions with characteristic scRNA-seq noise, including Gaussian noise and expression dropouts, represented in **Fig. 1a**. The ground-truth network comprised 50 gene pairs, each assigned one of the four interaction types (**Supplementary Fig. 1**). Using simulated data and ground-truth networks, we first evaluated the over-all root-mean-square error (RMSE) for algorithms that bound gene dependencies within the [0, 1] interval, analogous to traditional correlation coefficients. Chatterjee’s correlation coefficient outperformed the Pearson, Spearman, and Distance correlation metrics, yielding the lowest overall RMSE (**Fig. 1b**). ChatterjeeKNN, the k-nearest-neighbor variant of the coefficient, reduced the RMSE further.

**Fig 1.**
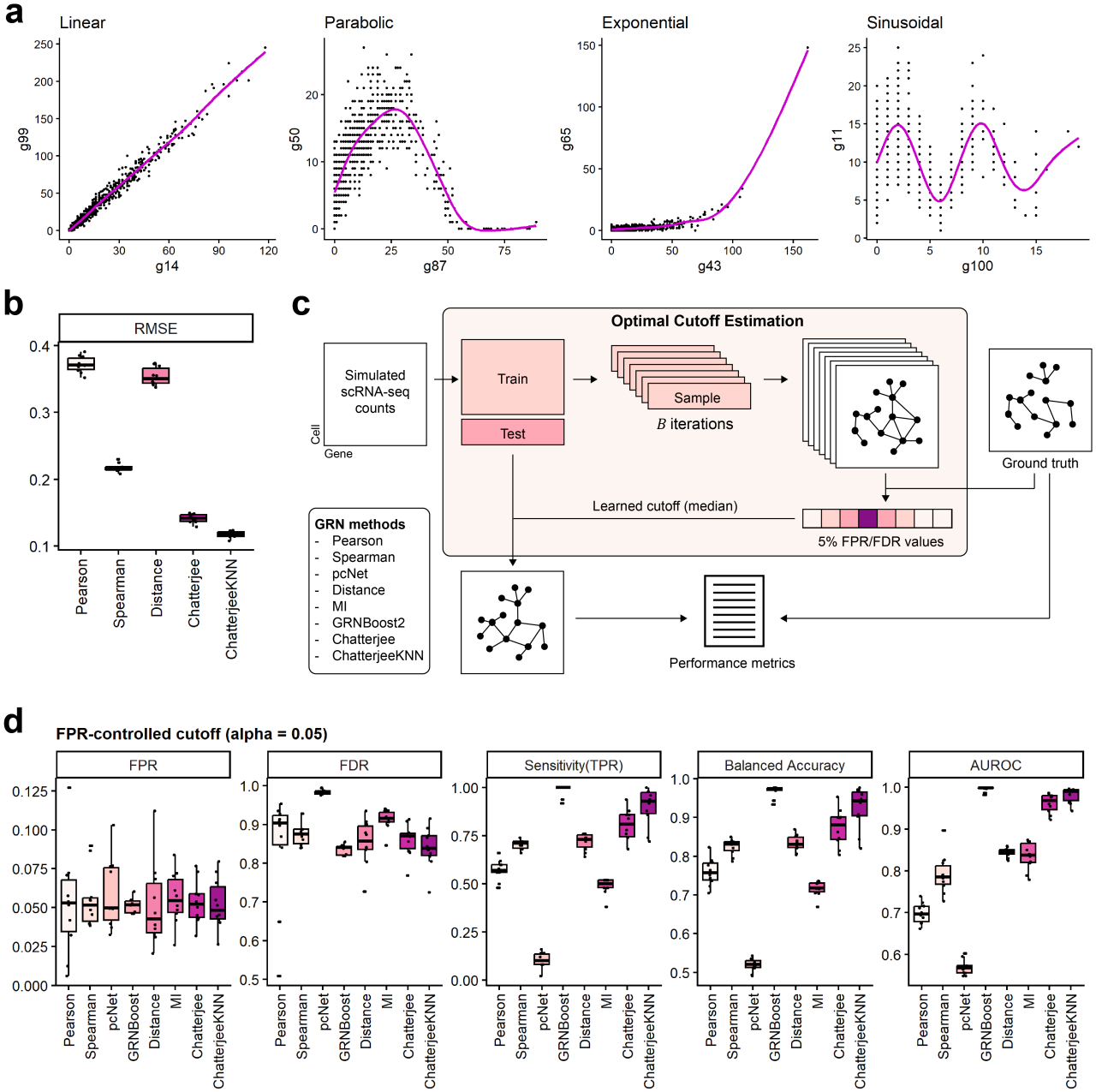
Evaluation of Chatterjee’s correlation and performance benchmarking on simulated scRNA-seq data with non-linear gene dependencies. (a) Representative scatter plots showing four distinct simulated gene-gene expression relationships: linear, parabolic, exponential, and sinusoidal. (b) Comparison of RMSE for methods constraining gene dependency scores within the [0, 1] interval. (c) Schematic overview of the optimal cutoff selection framework. Training counts (*n*_train_ = 600) are iteratively sub-sampled (*B* = 50) against ground-truth links to learn empirical cutoffs that strictly control the false positive rate (FPR) at *α* = 0.05 before application to test data (*n*_test_ = 400). (d) Performance of eight GRN inference methods across test datasets using the following classification metrics: FPR, false discovery rate (FDR), sensitivity (TPR), balanced accuracy, and AUROC, with cutoffs controlling FPR at *α* = 0.05.

To broaden our benchmarking, we next compared Chatterjee’s coefficient against established GRN inference tools designed for RNA-seq data, specifically pcRegression (pcNet), mutual information (MI), and the random forest-based GRNBoost2. Because these methods yield scale-invariant or unbounded metrics that can approach infinity, they generate widely disparate score distributions that don’t allow straight-forward RMSE-based comparisons (**Supplementary Fig. 2b**). To accommodate these unbounded metrics, we reframed the benchmark as a classification problem, which aligns with the primary objective of identifying discrete gene dependencies, and employed a data-driven non-parametric optimal cutoff estimation approach (Figure 1c). To estimate thresholds that strictly control the false positive rate (FPR) at 0.05 across all algorithms, each dataset was randomly partitioned into training (*n*_train_ = 600 cells) and test (*n*_test_ = 400 cells) sets. We then applied Algorithm 1 to the training data (setting *n*′ = *n*_train_, *n* = *n*_test_, *B* = 50, and *α* = 0.05) to derive empirical cutoffs. These cutoffs were subsequently applied to the testing procedures detailed in (7) and (8), ensuring uniform FPR control when evaluating the test data.

**Fig 2.**
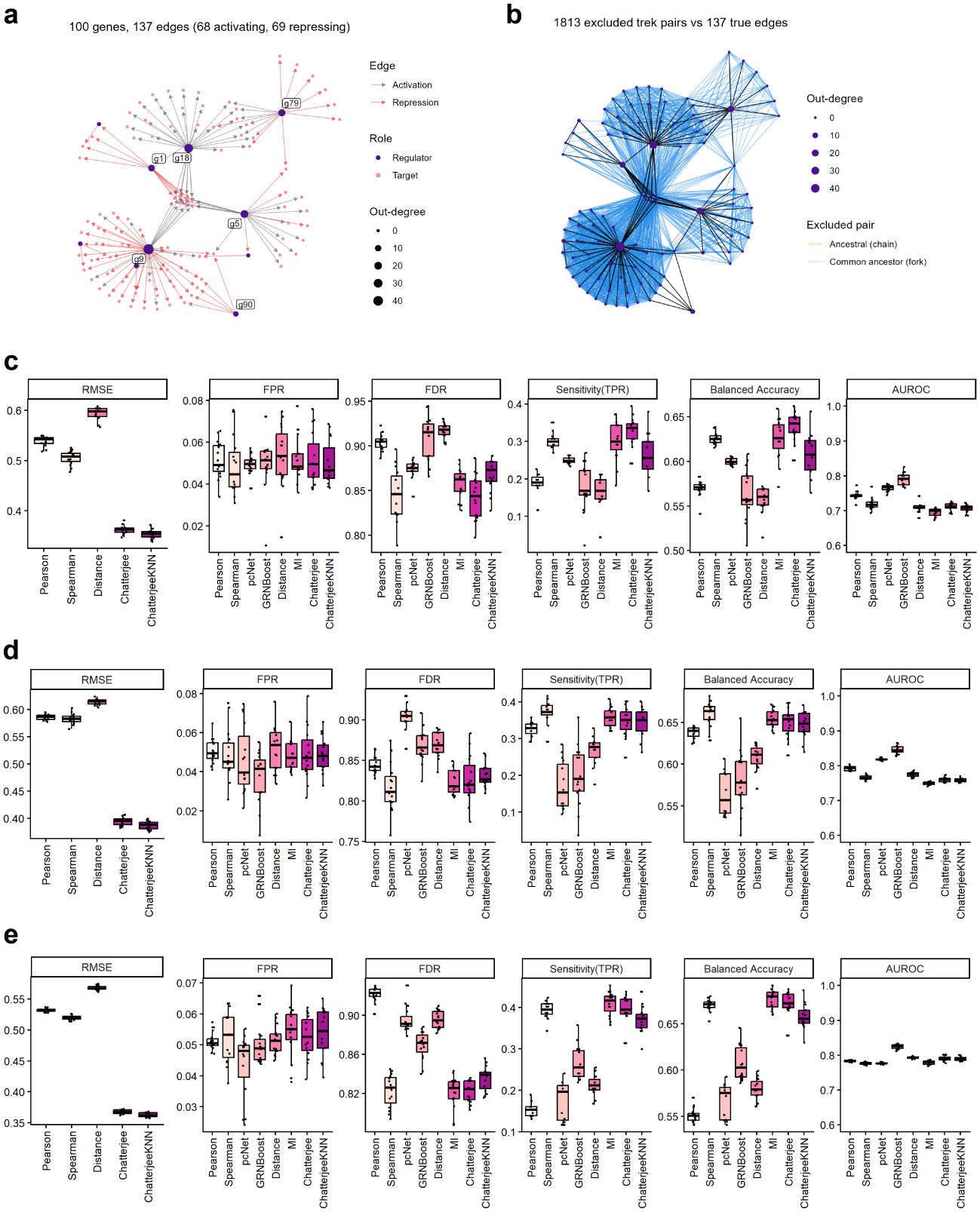
Performance benchmarking and GRN inference evaluation across diverse SERGIO-simulated data. (a) Ground-truth GRN shared by DS4, DS5, and DS6 (100 genes, 137 edges: 68 activating, 69 repressing). Node size indicates out-degree, regulators are shown in purple, and master regulators are labeled. (b) Gene pairs connected by a trek (ancestral chain or common-ancestor fork) but not directly linked: 1,813 pairs versus 137 true edges. True edges are shown in black, and excluded pairs in orange (chain) and blue (fork). These pairs are excluded from the null set only in the analysis of **Supplementary Fig. 5**. (c–e) Benchmarking results for DS4 (c), DS5 (d), and DS6 (e), with all non-edges treated as nulls. In each row, the first panel shows RMSE for the [0, 1] bounded dependency metrics (Pearson, Spearman, Distance, Chatterjee, and ChatterjeeKNN correlation). The remaining panels show FPR, FDR, sensitivity (TPR), balanced accuracy, and AUROC for eight network inference methods, with cutoffs estimated to control the FPR at 0.05.

Applying the estimated cutoffs to the test data demonstrated that our thresholding procedure effectively controlled the FPR at approximately 0.05 across all evaluated methods (**Fig. 1d**), establishing a rigorous baseline for comparative evaluation. Because true edges are sparse (50 of 4,950 gene pairs), the false discovery rate (FDR) at this threshold exceeded 0.8 for all methods (**Fig. 1d**).

In terms of sensitivity and balanced accuracy, Chatterjee’s correlation outper-formed all traditional correlation measures (Pearson, Spearman, and Distance), pcNet, and MI, while achieving competitive results with GRNBoost2 (**Fig. 1d** and **Supplementary Table 2**). ChatterjeeKNN further improved sensitivity and balanced accuracy over the standard coefficient, approaching GRNBoost2. The low sensitivity of pcNet in this setting is expected given its design: with its default of three principal components, pcNet assumes that gene-gene dependencies are driven by a few shared expression programs, whereas our simulation comprises 50 independent gene pairs whose covariance is spread across many dimensions and therefore poorly captured by the leading components. Consistent with this, pcNet performed competitively on the SERGIO datasets (Section 2.1.2), where regulatory structure is dominated by a small number of master regulators. Crucially, the area under the receiver operating characteristic (AUROC) curve of Chatterjee’s correlation was comparable to that of GRNBoost2, with performance discrepancies restricted to the order of 10^−2^ (**Fig. 1d**). To address the high FDR, we re-estimated the cutoffs with Algorithm 1, controlling the FDR instead of the FPR at 0.05. This reduced the FPR below 0.005 and the median FDR below 0.1 for all methods, except pcNet, which detected no edges (**Supplementary Fig. 2a**). Under this criterion, Chatterjee’s correlation and ChatterjeeKNN retained higher sensitivity and balanced accuracy than Pearson and MI and were comparable to Distance, while Spearman and GRNBoost2 were slightly higher (**Supplementary Fig. 2a**). The FDR-controlled cutoffs were stricter than the FPR-controlled cutoffs for every method (**Supplementary Fig. 2b**).

Finally, empirical runtime benchmarking (**Supplementary Fig. 3**) demonstrated that while Chatterjee’s correlation was marginally slower than Pearson, Spearman, and pcNet, it scaled more efficiently than distance correlation and MI as the dataset dimension increased. In single-core runs (**Supplementary Fig. 3a**), Chatterjee’s runtime remained low across all gene and cell counts, whereas distance correlation exceeded 10,000 seconds at 1,000 genes and 3,000 cells, and MI reached several hundred seconds. ChatterjeeKNN was slower than Chatterjee but remained faster than distance correlation. With 8 cores for distance correlation, Chatterjee, and ChatterjeeKNN (**Supplementary Fig. 3b**), distance correlation and single-threaded MI remained the slowest methods at 1,000 genes, while Chatterjee stayed at the low end of the range. Parallelization added overhead at small gene counts, so the multicore advantage is clearest for large datasets. The runtime of Chatterjee’s correlation therefore scales favorably with both genes and cells, supporting its use on large scRNA-seq datasets. Collectively, these findings demonstrate that the rank-based Chatterjee’s correlation offers a substantial improvement over traditional association metrics. It achieves performance on par with complex machine-learning architectures while remaining computationally efficient and mathematically transparent.

**Fig 3.**
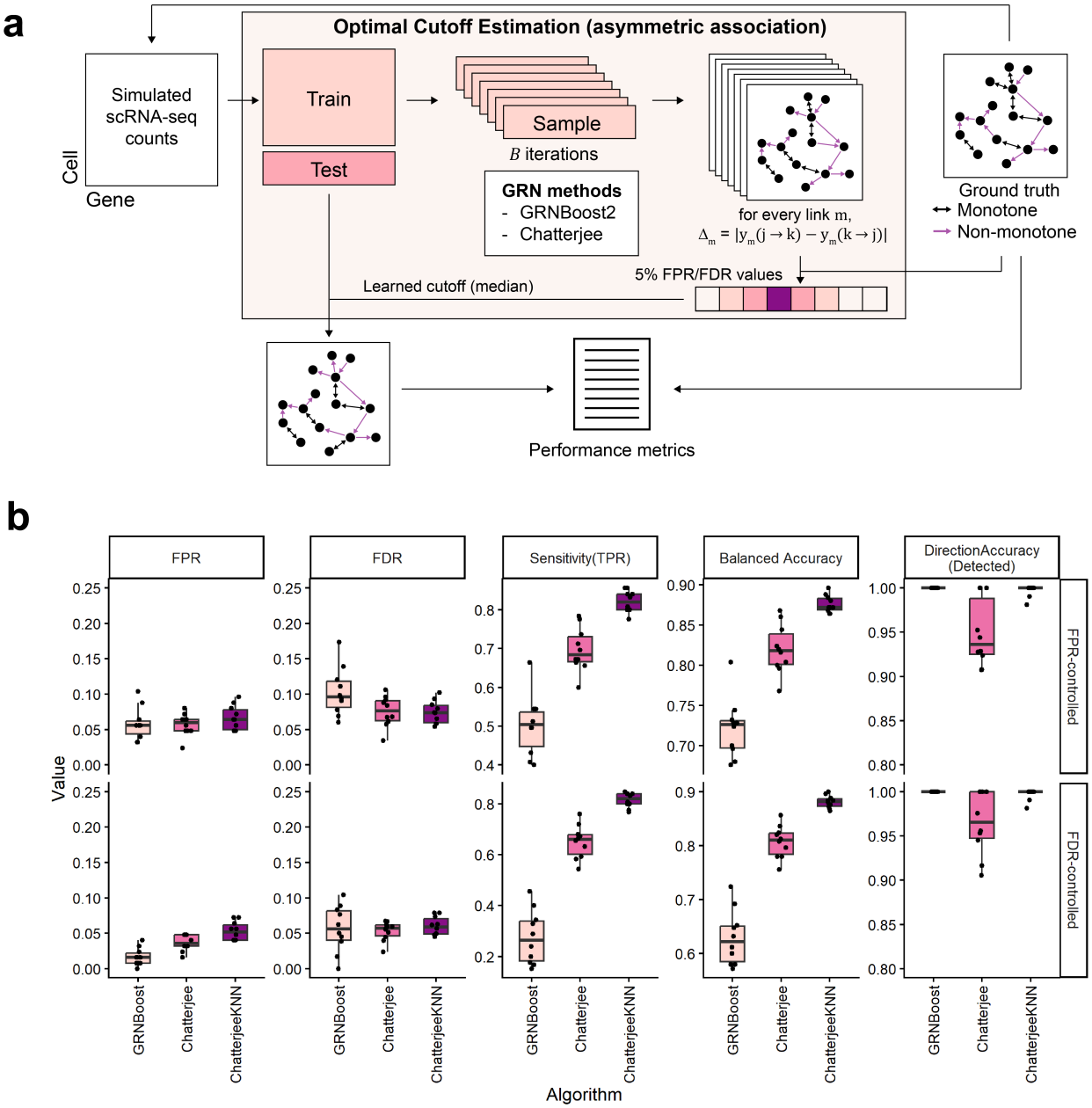
Chatterjee’s correlation coefficient identifies asymmetric predictive associations in simulated gene regulatory networks. (a) Schematic overview of cutoff estimation and evaluation of asymmetric association detection. Simulated scRNA-seq count data with known ground-truth non-monotone (asymmetric) and monotone (symmetric) gene-gene interactions were split into training and test sets. Using the training set, cutoffs on the directed dependency score, 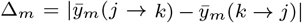, were learned over B resampling iterations to control either the FPR or the FDR at 5% (median across iterations) and then applied to the held-out test set. (b) Performance of network inference for GRNBoost2, Chatterjee, and ChatterjeeKNN, measured by FPR, FDR, sensitivity, balanced accuracy, and direction accuracy for detected asymmetric associations, under FPR-controlled (top row) and FDR-controlled (bottom row) cutoffs.

#### 2.1.2 Second study: Benchmarks on SERGIO simulated data

To evaluate performance on more complex and biologically realistic topologies, we utilized the scRNA-seq simulator SERGIO [23]. SERGIO models cell differentiation trajectories characteristic of real scRNA-seq data using user-defined GRNs, which provide a definitive ground-truth network structure necessary for optimal cutoff selection and validation. The three datasets used here, DS4, DS5, and DS6, share a single ground-truth GRN of 100 genes and 137 regulatory edges (68 activating, 69 repressing), structured as a shallow DAG driven by six master regulators (**Fig. 2a**). This network is sparse and hub-dominated: 62% of genes carry a single edge, whereas five master regulators have degree ≥ 12 (maximum 47), yielding a heavy-tailed degree distribution (**Supplementary Fig. 5a**). The datasets differ only in their differentiation graphs, which are linear (DS4, 3 cell types), bifurcating (DS5, 4 cell types), and trifurcating (DS6, 6 cell types), and in their cell type-specific master regulator production rates (**Supplementary Fig. 4**). Holding the network topology fixed therefore isolates the effect of trajectory structure on GRN inference. In this section, we benchmark GRN inference algorithms across these three SERGIO-simulated datasets and their distinct cell trajectory structures (**Supplementary Fig. 4**a, d, g). Adopting the benchmarking workflow established in our non-linear simulations, we first applied Algorithm 1 to determine empirical cutoffs for each method within each dataset, followed by a rigorous comparative performance analysis.

**Fig 4.**
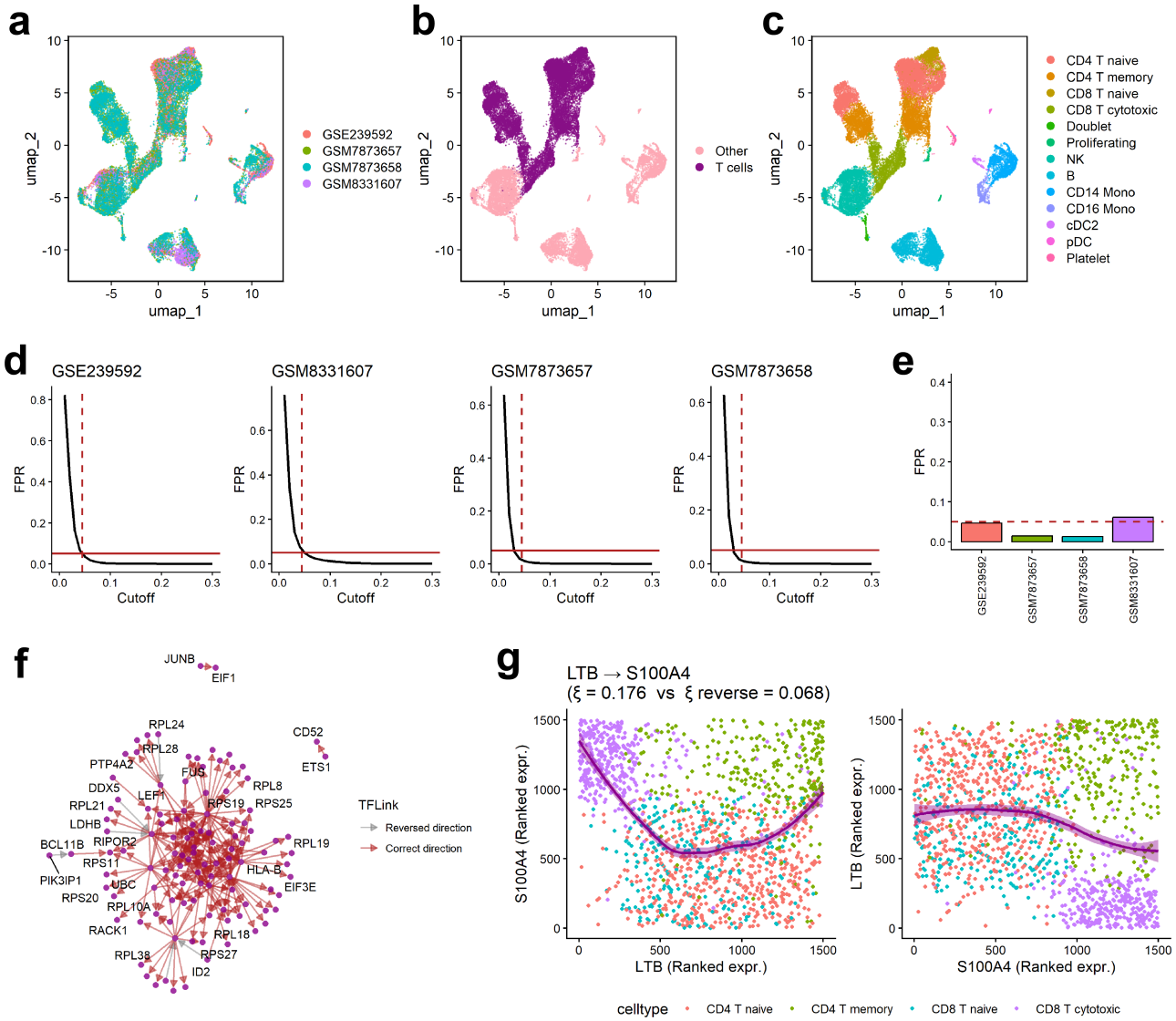
GRN construction from real PBMC scRNA-seq data with asymmetric associations. (a) UMAP embedding of four integrated PBMC datasets (GSE239592, GSM8331607, GSM7873657, GSM7873658) after Harmony batch correction. (b) Same UMAP colored by cell-type classification (T cells vs. Other), used to isolate the T-cell population for downstream analysis. (c) Same UMAP colored by annotated PBMC cell types. (d) FPR as a function of the dependency-testing cutoff for each of the four PBMC datasets. The cutoff was estimated once from the GSE239592 training set via Algorithm 1 (dashed vertical line) and applied unmodified to the three held-out datasets. The solid horizontal line marks the target FPR of 0.05. (e) Realized FPR in each of the four datasets when the GSE239592-derived cutoff is applied; the dashed line marks 0.05. (f) Directed network of significantly asymmetrically associated gene pairs identified in the GSM8331607 dataset using the asymmetric association test, restricted to pairs annotated in TFLink; arrows point from predictor to target gene. Red edges match the TFLink-annotated direction (transcription factor to target), and gray edges are reversed. (g) Asymmetry underlying the inferred direction, depicted by the ranked-expression relationship between LTB and S100A4, the top-ranked asymmetric association in GSM8331607 (Supplementary Fig. 9b), with cells colored by T-cell subtype.

Chatterjee’s coefficient consistently achieved the lowest RMSE compared to the other bounded [0, 1] metrics—Pearson, Spearman, and Distance correlation—across all three SERGIO datasets (Figure 2c–e). ChatterjeeKNN reduced the RMSE further in all three datasets. For the downstream binary classification tasks, our empirical cutoff estimation framework successfully constrained the FPR to approximately 0.05 across all evaluated methods, establishing a uniform baseline for unbiased performance comparison (Figure 2c–e).

A comprehensive evaluation across these trajectories (**Supplementary Tables 3, 5, 7**) revealed that Chatterjee’s correlation achieved the highest sensitivity and balanced accuracy in DS4, while performing comparably to top-performing methods in DS5 and DS6. In DS5, traditional metrics such as Pearson and Spearman correlation also performed robustly. Notably, while GRNBoost2 consistently yielded high AUROC scores, it exhibited low balanced accuracy under the fixed 0.05 FPR threshold. This disparity suggests that although random forest-based approaches excel at global edge-ranking, their performance degrades at strict, uniform FPR cutoffs when applied to highly imbalanced network datasets where true regulatory links are sparse relative to non-interacting pairs. Consequently, while GRNBoost2 serves as an effective ranking tool, it lacks a theoretically grounded framework for stable threshold determination.

Despite the controlled FPR, the FDR remained high for all methods (approxi-mately 0.8–0.9; **Fig. 2c–e**), as expected for a network with 137 true edges among 4,950 gene pairs. We could not obtain FDR-controlled cutoffs on these data using Algorithm 1 with the full set of non-edges as the null. In SERGIO, many non-edges are dependent through shared upstream regulators, and these indirect dependencies inflate the null score distributions. To address this, we excluded trek pairs, i.e., non-adjacent gene pairs connected by a directed path (chain) or sharing a common ancestor (fork), from the null set. This removed 1,813 pairs, compared with 137 true edges (**Fig. 2b**). With trek pairs excluded, FDR-controlled cutoffs lowered the FPR to below 0.02 and the median FDR to below 0.1 for most methods in all three datasets (**Supplementary Fig. 5e, g, i**; **Supplementary Tables 4, 6, 8**), whereas FPR-controlled cutoffs continued to yield FDRs of 0.5–0.7 (**Supplementary Fig. 5d, f, h**). Under FDR control, Chatterjee’s correlation and ChatterjeeKNN were comparable to Pearson, Spearman, and Distance correlation in DS5 and DS6 and outperformed MI and pcNet in sensitivity and balanced accuracy, whereas Pearson, Spearman, and Distance correlation were higher in DS4 (**Supplementary Fig. 5e, g, i**). The RMSE ranking was unchanged, with ChatterjeeKNN and Chatterjee lowest (**Supplementary Fig. 5c**).

When paired with our optimal cutoff estimation framework, Chatterjee’s coefficient emerges as a robust alternative. Although it exhibits slightly lower global AUROC than learning-based methods, indicating weaker overall edge ranking, our data-driven thresholding strategy effectively resolves this limitation for Chatterjee’s correlation in complex simulations. Collectively, these results demonstrate that Chatterjee’s correlation substantially outperforms traditional correlation measures and remains highly competitive with advanced non-parametric metrics like Distance correlation and MI. The two variants performed similarly overall. ChatterjeeKNN had the lowest RMSE in all three SERGIO datasets and matched or slightly exceeded Chatterjee’s sensitivity under FDR control with trek pairs excluded in DS4 and DS6 (**Supplementary Fig. 5e, i**), whereas the standard coefficient had higher sensitivity and balanced accuracy under FPR-controlled cutoffs in DS4 and DS6 (**Fig. 2c, e**).

In summary, benchmarks across both simulation cases demonstrate that our proposed multiple-testing framework utilizing Chatterjee’s correlation coefficient significantly outperforms traditional association measures in identifying dependent gene pairs and reconstructing underlying GRNs from scRNA-seq data. Furthermore, this metric provides robust classification accuracy at a lower computational cost than complex, learning-based approaches. While these results establish the efficacy of Chatterjee’s correlation as a symmetric association framework, a major advantage of this metric lies in its inherent and interpretable statistical asymmetry. Consequently, the next section explores the directional properties of Chatterjee’s correlation to infer asymmetric predictive associations within GRNs.

### 2.2 Chatterjee’s correlation accurately identifies asymmetric predictive associations in simulated scRNA-seq data

Chatterjee’s correlation coefficient has a distinctive property that under non-linear dependencies it is non-symmetric, i.e., *ξ*(*i* → *j*) ≠ *ξ*(*j* → *i*), and this asymmetry vanishes under strictly linear relationships. This offers a natural, rank-based means of inferring regulatory direction directly from a pairwise statistic, without requiring a global model of the network. In this section, we test whether this asymmetry, defined as a significant difference between *ξ*(*i* → *j*) and *ξ*(*j* → *i*), can reliably identify asymmetrically associated (directed) gene pairs, using the testing procedure in (10). GRNBoost2 similarly produces an asymmetric, directed importance score for each gene pair (11), so we included it as a comparator to evaluate whether Chatterjee’s simple pairwise formulation could match or exceed the performance of this more complex, tree-ensemble-based approach.

To this end, we simulated synthetic scRNA-seq count data with predefined genegene interactions, following the approach described in Section 3.4. Each simulated dataset comprised 500 genes and 2000 cells with a 20% dropout rate, and included 250 labeled gene-gene interactions classified as either monotone (symmetric) or non-monotone (asymmetric) (**Fig. 3a**).

For each simulated dataset, cells were randomly partitioned into a training set (*n*_train_ = 1200) and a test set (*n*_test_ = 800). Following Algorithm 2, the training set was used to determine testing cutoffs for both procedures (10) and (11), with *n*′ = *n*_train_, *n* = *n*_test_, *B* = 50, and *α* = 0.05, based on the known ground-truth interactions from the simulation. From the same resamples, we additionally estimated a cutoff controlling the false discovery rate (FDR) at *α*, defined as the threshold retaining the largest set of top-ranked gene pairs whose empirical false discovery proportion did not exceed *α*. The independent test set was reserved for subsequent performance evaluation.

We applied the estimated cutoffs to the asymmetric GRNs reconstructed from the test set using both GRNBoost2 and Chatterjee’s correlation, as well as its k-nearest-neighbor variant ChatterjeeKNN (*M* = 5), classifying a gene pair as asymmetrically associated whenever the asymmetry in its dependency measure exceeded the corresponding cutoff. Under the FPR-controlled cutoffs, all methods controlled the FPR near the target level of 0.05 (**Fig. 3b**, top row; **Supplementary Table 9**). Chatterjee’s correlation identified substantially more true asymmetric associations than GRNBoost2, with a higher median sensitivity and correspondingly higher balanced accuracy, as well as a lower realized FDR. ChatterjeeKNN further increased sensitivity (≈ 0.82) and balanced accuracy (≈ 0.88) relative to Chatterjee’s correlation (≈ 0.69 and ≈ 0.82).

Under the FDR-controlled cutoffs, all methods kept the realized FDR near 0.05 (**Fig. 3b**, bottom row). However, while the sensitivity of Chatterjee’s correlation and ChatterjeeKNN was largely unchanged (≈ 0.66 and ≈ 0.82 vs. ≈ 0.69 and ≈ 0.82), GRNBoost2’s sensitivity dropped by roughly half (≈ 0.26 vs. ≈ 0.50). This suggests that true asymmetric associations are concentrated at the top of the asymmetry ranking of Chatterjee’s correlation, whereas in GRNBoost2 they are interleaved with false ones.

The learned cutoffs were stable with sample size. The cutoff decreased smoothly with the number of cells *n* and followed the fitted power-law function under both FPR and FDR control (**Supplementary Fig. 6a**). When applied to held-out cells, the realized FPR and FDR approached the 0.05 target as *n* increased, and sensitivity plateaued at approximately 0.83 beyond roughly 1,800 cells, with the largest variability at small *n* (**Supplementary Fig. 6b**).

In contrast, the cutoffs did not transfer across simulation settings (**Supplementary Fig. 7**). Cutoffs learned with Algorithm 2 on data matching the test setting kept the realized FPR between 0.03 and 0.07 (**Supplementary Fig. 7b**, diagonal), whereas the baseline formula failed under large noise (noise standard deviation 1.5, FPR ≈ 0.15–0.30) and high dropout (dropout rate 0.5, FPR ≈ 0.12–0.22), with the error growing with *n* (**Supplementary Fig. 7a**). Transfer was asymmetric. Cutoffs learned at low dropout (0–0.2) produced FPRs of up to 0.44 when applied to higher-dropout data, whereas cutoffs learned at high dropout were conservative (FPR ≤ 0.09) in the other settings. The cutoff therefore needs to be relearned when the data characteristics change, which motivates estimating it from a reference dataset for real data (Section 2.3).

When restricted to pairs already correctly labeled as asymmetrically associated, GRNBoost2 and ChatterjeeKNN assigned the correct direction almost without exception (median ≈ 1.0), and Chatterjee’s correlation did so with slightly lower accuracy (median ≈ 0.94–0.97), under either cutoff. This indicates that once a regulatory link is detected, all methods reliably recover its direction. The principal advantage of the Chatterjee coefficients lies instead in their higher sensitivity for detecting asymmetric associations in the first place, achieved through a simple, pairwise, rank-based test rather than GRNBoost2’s ensemble-based inference over the full network structure, an advantage that widens under FDR control. Given the higher sensitivity, balanced accuracy, and direction accuracy of ChatterjeeKNN, combined with its interpretability and computational efficiency, we adopt ChatterjeeKNN (*M* = 5) as our primary method for testing asymmetric predictive associations in the real-data analyses that follow.

Accordingly, in Section 2.3 we apply the testing procedures (7) and (10), based on ChatterjeeKNN, to a real scRNA-seq dataset to recover the underlying gene regulatory network and biologically validate the inferred regulatory links.

### 2.3 Directed GRN construction from real data using Chatterjee’s correlation

Building on the simulation results in Section 2.2, which showed that Chatterjee’s correlation outperforms GRNBoost2 in identifying non-monotone (asymmetric) interactions, we now turn to real single-cell data to identify gene pairs with asymmetric association, as formalized in (9). Focusing on ChatterjeeKNN (*M* = 5), we apply the testing procedure in (10) with a cutoff learned on the data.

Given the dynamic, context-specific nature of biological systems, we evaluated the transferability of our data-driven cutoff selection procedure across independent PBMC datasets. Four scRNA-seq PBMC datasets (GSE239592, GSM8331607, GSM7873657, GSM7873658) were merged and integrated with Harmony (**Fig. 4a**), and T cells were identified from the annotated cell types (**Fig. 4b**,**c**; **Supplementary Fig. 8a**,**b**). Restricting to a common panel of genes detected in more than 30% of T cells in every dataset (**Supplementary Table 10**) and 1500 cells per dataset, we estimated a single dependency-testing cutoff on the GSE239592 dataset alone using Algorithm 1 (*B* = 50 resampling iterations, *α* = 0.05, with high-confidence (combined score *>*900) STRING [24, 25] interactions serving as ground truth). This cutoff, estimated once from GSE239592, was then applied without modification to the three remaining PBMC datasets. As shown in **Fig. 4d**, it controlled the FPR at approximately the target level of 0.05 both in the training dataset and in all three held-out datasets, and the realized FPRs for the held-out datasets, summarized in **Fig. 4e**, remained close to this threshold: below it in GSM7873657 and GSM7873658 (≈ 0.01) and slightly above it in GSM8331607 (≈ 0.06). The four datasets had broadly similar distributions of zero fraction, mean, standard deviation, and dispersion (**Supplementary Fig. 8c**), and in simulations a learned cutoff transferred reliably only between datasets with similar characteristics (**Supplementary Fig. 7**). These results confirm that a cutoff estimated from a single reference PBMC dataset generalizes reliably across independent donors and experimental batches of the same cell type.

We next applied the dependency test ((7) with cutoff computed on GSE239592), followed by the asymmetric predictive association test to the GSM8331607 dataset to recover asymmetric associations among its significantly dependent gene pairs. The cutoff for the asymmetric test was learned with Algorithm 2, which requires known dependent gene pairs, of which the null set is the subset expected to be dependent but not asymmetrically associated. We took these pairs from CORUM [26], a curated database of experimentally characterized mammalian protein complexes. Subunits of a stable complex are co-regulated and co-expressed, so they are dependent, but this co-expression is expected to be largely monotone and symmetric (e.g., HLA-B and B2M), so neither gene should predict the other asymmetrically. CORUM is therefore a suitable null for asymmetry, whereas STRING, which also contains regulatory and signaling links, is not. Because CORUM complexes are curated from low-throughput experiments, they also provide a high-confidence set. The learned cut-off was applied unchanged to GSM8331607. This yielded 18,232 dependent gene pairs (**Supplementary Fig. 9a**) and 5,134 gene pairs with significant asymmetric association (**Supplementary Fig. 9b**; pairwise relationships in **Supplementary Fig. 10**). To assess the inferred direction, we compared the asymmetric pairs with the transcription factor (TF)–target annotations in TFLink [27]. Of the 223 pairs annotated in TFLink, 217 (97.3%) had the inferred predictor-to-target direction matching the annotated TF-to-target direction, and the remainder were reversed (**Fig. 4f**).

The top-ranked asymmetric association, LTB → S100A4, as examined by its ranked expression relationship in **Fig. 4g**, showed a clear non-monotone and sub-stantially stronger relationship in the forward direction (*ξ* = 0.176) than in reverse (*ξ*_reverse_ = 0.068), illustrating how the asymmetry of Chatterjee’s coefficient resolves direction directly from expression data. LTB is an upstream activator of NF-*κ*B [28], a key transcriptional regulator of S100A4 [29]. Coloring cells by T-cell subtype showed that the non-monotone shape reflects subtype composition. CD4 memory T cells combined high LTB with high S100A4, consistent with LTB expression in CD4 memory T cells [30] and S100A4 enrichment in both CD4 memory and CD8+ cytotoxic T cells [31]. CD8 cytotoxic T cells had low LTB, whereas naive T cells showed moderately high LTB [32]. This pattern therefore likely reflects differences in CD4/CD8 lineage and memory/effector state rather than a direct regulatory mechanism. Chatterjee’s coefficient is a measure of statistical association, not causation, so an asymmetric association indicates that one gene predicts the other, not that it regulates it. Experimental validation of this interaction is beyond the scope of this study.

FDR could not be controlled on real data because high-confidence STRING interactions are sparse relative to non-interacting pairs and STRING is incomplete: genuine co-expression without a recorded protein association is counted as a false positive. We therefore calibrate and report FPR on real data, which depends only on the null distribution and transferred across datasets (**Fig. 4d**,**e**), and reserve FDR evaluation for simulations with complete ground truth.

## 3 Methods

### 3.1 Acquisition and pre-processing of real scRNA-seq data

Real scRNA-seq datasets were obtained from the Gene Expression Omnibus (GEO) database as detailed in **Supplementary Table 1**. The data was pre-processed using the Seurat(v5) R package [33]. For GSE239592, only control samples were used. Cells with greater than 500 detected genes, greater than 1000 reads, and less than 10% mito-chondrial reads were retained for subsequent analysis, and genes detected in fewer than 10 cells were removed. Doublets were identified in each dataset with scDblFinder [34] and removed. The four PBMC datasets were then merged. The data was then normalized using the “LogNormalize” function with a scale factor of 10,000. Highly variable features were identified using “FindVariableFeatures”, and the data was scaled using “ScaleData” using default parameters. Then, dimensionality reduction was performed by applying Principal Component Analysis (PCA) with 30 components, and datasets were integrated with Harmony using the Seurat v5 layer-aware interface. UMAP was computed on the first 20 Harmony dimensions for visualization. Finally, cells were clustered using the Louvain algorithm with a resolution of 1 on the Harmony embedding (20 dimensions, 20 nearest neighbors), and clusters were annotated into PBMC cell types from canonical marker genes (**Supplementary Fig. 8a**). For the Peripheral Blood Mononuclear Cells (PBMC) datasets only, T cell clusters (CD4 naive, CD4 memory, CD8 naive, and CD8 cytotoxic) were extracted based on these annotations, which relied on *CD3G, CD3E*, and *CD3D* expression among other markers. Genes detected in fewer than 30% of the cells in any dataset were then removed, and T cells with fewer than 10 total counts were dropped, leaving the common gene panel used in all analyses. Each pre-processed scRNA-seq dataset is referenced as the respective gene expression matrix in the following subsection.

### 3.2 Simulation of scRNA-seq data using SERGIO

To evaluate performance on more biologically realistic network topologies, we generated synthetic scRNA-seq data using SERGIO [23], which simulates gene expression along user-defined ground-truth GRNs and cell differentiation trajectories. We utilized three simulated datasets along with their respective ground truth, DS4, DS5, and DS6, using SERGIO’s dynamic simulation mode available in https://github.com/PayamDiba/SERGIO. Each dataset contains 100 genes and 300 cells per cell type (3, 4, and 6 cell types for DS4, DS5, and DS6), and all three share the same ground-truth GRN. Each dataset had 15 replicates, and we used the noise-free expression matrices. For each dataset, cells were partitioned into a training set (*n*_train_ = 60%) and a test set (*n*_test_ = 40%), and testing cutoffs for all eight GRN construction methods were estimated on the training set following Algorithm 1, with *B* = 50 and *α* = 0.05, as described in Section 2.1.2. Cutoffs were estimated to control either the FPR or the FDR at *α*. Gene pairs were labeled as true edges (direct regulation) or nulls (no trek connection in the ground-truth GRN). Pairs connected by a trek, i.e., sharing an ancestral relation or a common ancestor but not directly linked, were excluded from the null set and from evaluation (Supplementary Fig. 5); in the analysis without this exclusion, all non-edges were treated as nulls.

### 3.3 Simulation of scRNA-seq data with gene-gene interactions

We simulated scRNA-seq count data with predefined gene-gene interactions, including both linear and non-linear relationships. For each simulated dataset, we generated data for 100 genes and 1000 cells. Gene expression was modeled using a Poisson approximation.

For each gene *j* and cell *i*, a theoretical mean expression level was calculated as the product of a gene-specific mean and a cell-specific size factor. Gene-specific means were drawn from a Gamma distribution with shape parameter 1*/σ* and scale parameter *µ* × *σ*, where *µ* was set to 15, and *σ* was set to 0.5. Here *µ* and *σ* represent the mean expression and dispersion of gene *j*, respectively. Cell-specific size factors were drawn from a Gamma distribution with shape 1 and scale 1. To approximate Poisson-distributed counts, we used Bernoulli trials via the binomial distribution. The number of trials was set to a maximum of 1000, and ten times the theoretical mean expression, and the probability of success was set to the theoretical mean divided by the number of trials. This approach approximates a Poisson distribution. A dropout rate of 0.3 was introduced by randomly setting counts to zero with a probability of 0.3.

To introduce gene-gene interactions, we selected 50 gene pairs, each of which consists of a target gene and a source gene. The expression of the target gene was determined by the expression of the source gene as

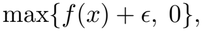

where *x* represents gene expression counts of the source gene, and *ϵ* represents a normally distributed noise variable with a standard deviation of 0.2. For the signal function *f* (*·*) we consider the following four forms:

1. **Linear**: *f* (*x*) = 2*x*
2. **Parabolic**: *f* (*x*) = −0.02 *x*^2^ + *x* + 5
3. **Exponential**: *f* (*x*) = *e*^5*x/θ*^, where *θ* = max expression of source gene
4. **Sinusoidal**: *f* (*x*) = 5 sin (15*x/θ*) + 10, where *θ* is the maximum expression of the source gene

After applying the gene relationship through the above functions, the resulting expression values for the target gene were converted back to count data using the same Poisson approximation method described above; no additional dropout was applied to target genes. The gene-gene interactions thus simulated are depicted in **Fig. 1a**. Ten independent datasets were simulated, each with a unique random seed. The 50 pairs are disjoint, so the remaining 4,900 gene pairs are independent and serve as the true nulls. Linear and exponential relationships were classified as monotone (symmetric), while parabolic and sinusoidal relationships were classified as non-monotone (asymmetric).

### 3.4 Simulation of scRNA-seq data with directed gene-gene interactions

To benchmark the asymmetric predictive association testing procedures described in Section 2.2 for Chatterjee’s coefficient, ChatterjeeKNN, and GRNBoost2, we applied the generative model from Section 3.3 to simulate scRNA-seq count data with a mixture of monotone and non-monotone (asymmetric) gene-gene interactions from Gamma-distributed gene means (*µ* = 15, *σ* = 0.5) and cell size factors, Bernoulli-approximated Poisson counts, and the same four signal functions (linear, parabolic, exponential, sinusoidal). Each simulated dataset comprised 500 genes and 2000 cells, with a dropout rate of 20%.

We defined 250 gene pairs, each consisting of a source and a target gene, by sampling 500 gene labels without replacement so that every gene participated in exactly one pair. The target gene’s expression was set as a function of the source gene’s expression plus Gaussian noise with standard deviation 0.2, cycling through the four signal functions in equal proportion and converting the resulting continuous values back to count data as before. Pairs generated under the linear and exponential functions were labeled monotone, and pairs generated under the parabolic and sinusoidal functions were labeled non-monotone. Ten independent replicate datasets were generated with distinct random seeds (1, …, 10), holding all other parameters fixed; within each replicate, cells were subsequently partitioned into training (*n*_train_ = 1200) and test (*n*_test_ = 800) sets as described in Section 2.2. Monotone pairs served as the null set for the asymmetry test.

### 3.5 Testing for dependence

#### 3.5.1 Multiple testing problem

We denote by N and ℝ the set of natural and real numbers, respectively, and for any *n* ∈ N, we denote [*n*] = *{*1, 2, …, *n}*. Suppose that we have *p* many distinct genes and for any *j* ∈ [*p*], the gene expression of the *j*^*th*^ gene is a random variable, denoted by *G*_*j*_. Due to the presence of biological regulation among genes, there is naturally an underlying dependence structure between the variables *G*_*j*_, *j* ∈ [*p*]. Thus, it is crucial to correctly identify these dependence relationships to construct a GRN. One of the simplest yet most efficient ways to comprehend the overall dependence structure is to study the degree of dependence in its fundamental units, i.e., gene pairs, and classify them as either dependent or independent. Therefore, one popular approach is to formulate a classification problem by considering multiple testing for pairwise independence among genes [35–38]. Formally, for every *j, k* ∈ [*p*] such that *j < k*, we consider the following test of hypothesis problem:

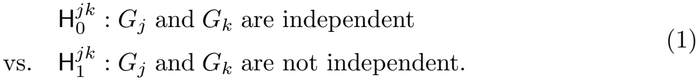

Furthermore, we consider the gene expressions of all *p* genes from *n* different cells which are basically *n possibly dependent* but identically distributed observations of the random vector *G* = (*G*_1_, *G*_2_, …, *G*_*p*_) ∈ ℝ^*p*^, given by *G*^(*i*)^ = (*G*_*i*1_, *G*_*i*2_, …, *G*_*ip*_), *i* ∈ [*n*]. These observations are stored as individual rows in a data matrix, also called the *gene expression matrix*, denoted by ***G*** ∈ ℝ^*n×p*^. Every (*i, j*)^th^ entry of ***G*** is represented by *G*_*ij*_, *i* ∈ [*n*], *j* ∈ [*p*].

For multiple testing correction, we used a false positive rate (FPR) based control. FPR is the expected ratio of the number of false positives to the total number of true null hypotheses, that is,

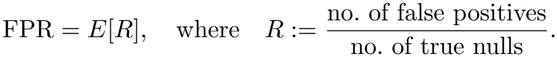

We also considered false discovery rate (FDR) based control. FDR is the expected ratio of the number of false positives to the total number of rejected null hypotheses, that is,

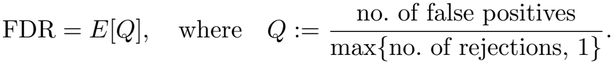

In all the following sections, both the FPR and the FDR for the above multiple testing problem (1), as well as for the asymmetric predictive association testing problem (10), were controlled at 0.05, with separate cutoffs estimated for each criterion in the simulations. On real data, only the FPR was controlled, since FDR could not be reliably evaluated without a complete ground truth (Section 2.3).

#### 3.5.2 Chatterjee’s rank-based correlation coefficient

There are numerous existing measures of dependence which can be implemented as test statistics for the multiple testing problem formulated in (1). In this work, our main focus is to propose a testing procedure based on *Chatterjee’s rank-based correlation coefficient* [11] to measure gene-gene dependencies, which we formalize as follows.

For any arbitrary *j, k* ∈ [*p*], we consider the *n* bivariate (paired) gene expression data (*G*_*ij*_, *G*_*ik*_), *i* ∈ [*n*] observed from the *j*^th^ and *k*^th^ genes. First, we rearrange the observations as

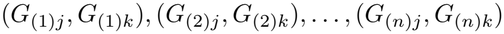

such that the observations of the *j*^th^ gene are ordered in an increasing fashion as

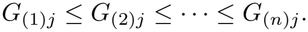

If *G*_*ij*_ ‘s and *G*_*ik*_’s have no ties, then there is a unique way to order the above. Subsequently, for every *i* ∈ [*n*], let *r*_*i*_ be the rank of *G*_(*i*)*k*_, i.e., *r*_*i*_ := |{*l* : *G*_(*l*)*k*_ ≤ *G*_(*i*)*k*_}|. Then, Chatterjee’s correlation coefficient is defined as, see [11, Section 1],

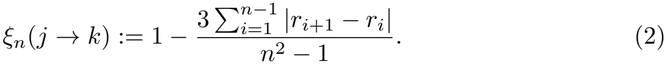

In the presence of ties, which is common in gene expression count data generated by scRNA-seq, the above coefficient is defined as follows. If there are ties among the *G*_*ij*_’s, then an increasing rearrangement as above is chosen by breaking the ties uniformly at random. Furthermore, regarding the *G*_*ik*_’s (with the possibility of ties), we additionally define the rank-type variable *l*_*i*_’s as *l*_*i*_ := |{*l* : *G*_(*l*)*k*_ ≥ *G*_(*i*)*k*_}|, and define the coefficient as

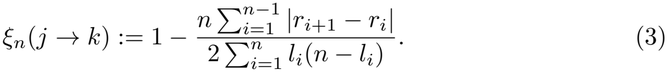

It is important to note that, since we first order *G*_*ij*_’s and then based on that assign the indices (*i*)*k*’s to *G*_*ik*_’s and subsequently obtain the ranks of *G*_(*i*)*k*_’s denoted as *r*_*i*_’s, *ξ*_*n*_(*j* → *k*) is not symmetric in *j* and *k*. If *j* and *k* are swapped in the above operations, that is, start by ordering *G*_*ik*_’s, then the resulting coefficient is denoted as *ξ*_*n*_(*k* → *j*). Thus, to obtain a symmetric coefficient, we compute the statistic by considering the maximum of *ξ*_*n*_(*j* → *k*) and *ξ*_*n*_(*k* → *j*), and denote it as *ξ*_*n*_(*j, k*), i.e.,

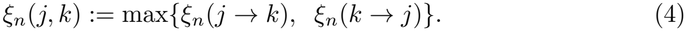

The above coefficient provides us with a measure of dependence between the variables *G*_*j*_ and *G*_*k*_, thereby estimating the degree of association or regulation between the *j*^th^ and *k*^th^ genes. In an asymptotic sense, when the sample size *n* is large enough and the cells are independent and identically distributed (IID), then *ξ*_*n*_(*j, k*) is close to 0 if the *j*^th^ and *k*^th^ genes are independent, i.e., there are no regulations between them. Conversely, *ξ*_*n*_(*j, k*) is closer to 1 in the presence of regulation [11, Theorem 1.1]. The strength of regulation is associated with how close *ξ*_*n*_(*j, k*) is to 1, and therefore it not only serves as an estimate of the dependence, but is also useful as a test statistic for testing of independence, stated in problem (1), which we describe in the next section.

Lin and Han [12] showed that the power of *ξ*_*n*_ can be boosted by using several right nearest neighbors in the rank construction instead of only the adjacent one. Keeping the ordering and ranks *r*_*i*_ defined above, and for a fixed integer *M* ≥ 1, the revised coefficient is

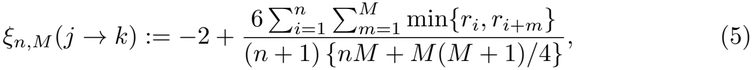

with the convention *r*_*i*+*m*_ := *r*_*i*_ whenever *i* + *m > n*. Here *G*_(*i*+*m*)*j*_ is the *m*^th^ right nearest neighbor of *G*_(*i*)*j*_, and ties in both genes are broken uniformly at random, so the ranks *r*_*i*_ here are computed with random tie-breaking rather than maximum ranks. For *M* = 1, *ξ*_*n*,*M*_ is asymptotically equivalent to *ξ*_*n*_. If *M* grows with *n* (with *M/n* → 0), it estimates the same population measure while gaining power against weak dependence [12]. Like *ξ*_*n*_, it is asymmetric in *j* and *k*, so we symmetrize it in the same way as (4):

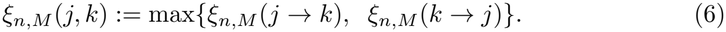

To distinguish it from the original coefficient, we refer to *ξ*_*n*,*M*_ as *ChatterjeeKNN* throughout this manuscript. Unless stated otherwise, we use *M* = 5.

We selected *M* on simulated data with known ground truth, before any real-data analysis. We evaluated *M* ∈ {1, 3, 5, 10, 20, 50} on the non-linear simulation of Section 3.3 (1,000 cells) and on the SERGIO datasets DS4, DS5, and DS6, using the first five replicates of each. Each replicate was split 60%/40% into training and test sets as in the benchmarks. For each *M*, we computed the symmetrized scores *ξ*_*n*,*M*_ (*j, k*) on the test set and measured the AUROC and the RMSE against the ground-truth labels over unique labeled gene pairs, which do not depend on a cutoff and therefore isolate the effect of *M* (**Supplementary Fig. 5b**). AUROC was nearly flat for *M* ≤ 10 in all four datasets (changes below 0.015) and declined for larger *M*, most strongly in the non-linear simulation (from approximately 0.99 at *M* = 10 to 0.92 at *M* = 50). In the non-linear simulation, RMSE was lowest at *M* = 5. In the SERGIO datasets, RMSE decreased steadily with *M*, but AUROC did not improve, so this reflects a shift in the score scale rather than better discrimination. We therefore chose *M* = 5, which stays within 0.01 of the best AUROC in every dataset (and within 0.005 for the non-linear simulation, DS5 and DS6) and gives the lowest RMSE in the non-linear simulation, while retaining more than one neighbor, which is the source of the power gain over *ξ*_*n*_ [12]. Because *M* ≤ 10 performed similarly, results are not sensitive to this choice within that range.

#### 3.5.3 Testing procedure

It has been shown in [11, Theorem 2.1] that, when *G*_*j*_ and *G*_*k*_ are independent, i.e., 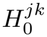 is true, and the observations from the cells are IID, then both 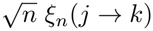 and 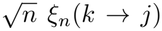 are asymptotically distributed as Normal with mean 0 and variance 2*/*5, i.e.,

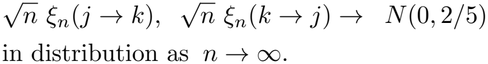

This result requires a continuous response and IID observations; it is not established for tied count data, for dependent cells, or for the symmetrized statistic *ξ*_*n*_(*j, k*), and we use it only as motivation for estimating the cutoffs empirically. It is noteworthy that the limiting distribution is not only symmetric but also independent of the true underlying distributions of *G*_*j*_ and *G*_*k*_, which are unknown. Thus, the limiting distribution of the symmetrized statistic 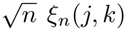 must also adopt such a favorable property. However, even under the assumption that the cell observations are IID, the specific parameter values are yet to be characterized in the statistical literature.

Nevertheless, in the spirit of the above results, to leverage the transparency and the distribution-free nature of the test statistics, we propose the testing procedure that,

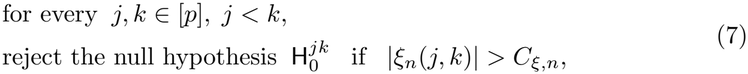

where *C*_*ξ*,*n*_ is some positive testing cutoff which must be determined appropri-ately, such that the FPR (or, alternatively, the FDR) is controlled by the required level. However, since any rigorous characterization of the limiting behavior or at least the limiting variance of *ξ*_*n*_(*j, k*) is yet absent in the existing literature, and more importantly, the cell observations are quite possibly dependent in practice, we need to estimate *C*_*ξ*,*n*_ separately from synthetic or pre-existing gene expression databases, where it is a priori known which gene pairs are independent. Subsequently, we use the estimated cutoff to apply the testing procedures on real datasets to identify the dependent gene pairs, as shown in Section 2.3.

In a similar way as in (7), we adopt a general testing policy which can incorporate the other dependency measures, resulting in the seven testing procedures listed below,

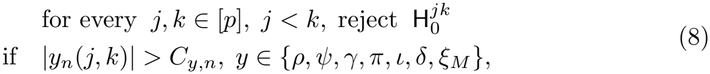

where the testing cutoffs *C*_*y*,*n*_, *y* ∈ {*ρ, ψ, γ, π, ι, δ, ξ*_*M*_}, similarly to *C*_*ξ*,*n*_, depend on the number of observations *n* and the corresponding limiting distributions of the coefficients. Although for the traditional measures, there are relevant asymptotic theories regarding their limiting distributions, their validity primarily depends on the crucial assumption that the cell observations must be IID, which does not hold in practice. Therefore, instead of using asymptotics or theoretical results, these testing cutoffs are similarly estimated from data, not only to avoid any error due to asymptotic approximations but also to perform a fair and more precise comparison with the procedure proposed in (7), as shown later in Section 2.1.

#### 3.5.4 Other dependency measures for GRN construction

We considered six additional dependency measures for GRN construction: two classical correlation coefficients (Pearson and Spearman), two nonparametric measures capable of capturing non-linear associations (distance correlation [6] and mutual information [7]), and two machine-learning-based network inference methods (GRNBoost2 [8] and PCNet [9]). All of the above measures were implemented to solve our testing problem (1), and their performances are discussed in the subsequent sections.

##### Pearson’s correlation coefficient

Pearson’s correlation coefficient is defined as

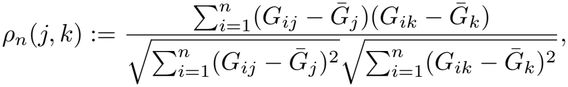

where 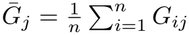 and 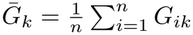.

##### Spearman’s rank correlation coefficient

First, for every *i* ∈ [*n*], we denote by *r*_*ij*_ and *r*_*ik*_ the ranks of *G*_*ij*_ and *G*_*ik*_ among their corresponding set of observations, respectively, i.e.,

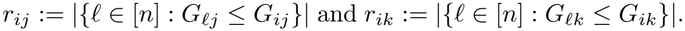

In the presence of ties, instead of assigning them consecutive ranks, we assign them the average of the ranks they would have otherwise received. Then, Spearman’s correlation coefficient is defined as

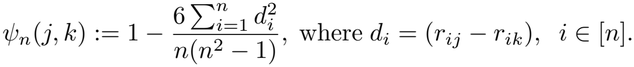

To account for global and multivariate relationships beyond simple pairwise statistics, we also included two ML-based dependency measures designed for network-wide GRN reconstruction.

##### GRNBoost2

In the case of GRNBoost2, we consider the dependency measure,

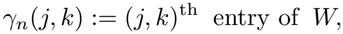

where *W* is the *symmetrized weight matrix* associated with the gene regulatory network that is constructed using the tree-based ensemble methods, as specified i n [8]. We used the arboreto Python implementation with default parameters, with all genes allowed as regulators. Absolute importance scores were arranged into a gene-by-gene matrix (zero for gene pairs not returned), and *W* was obtained by taking the maximum of the (*j, k*) and (*k, j*) entries, with the diagonal set to 0. For the asymmetric association test (11), the unsymmetrized importances were used, 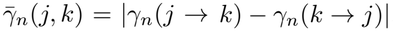.

##### PCNet

In the case of PCNet, we consider the dependency measure,

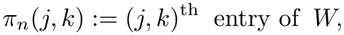

where *W* is the *symmetrized adjacency matrix* derived in the network construction and manifold alignment steps, as described in [9], using the default of three principal components.

***Distance correlation***

Distance correlation [6] is a measure of dependence that, unlike Pearson’s and Spear-man’s but similar to Chatterjee’s coefficients, is ca pable of de tecting non-linear associations and returns 0 if and only if the two variables are independent. For every *j, k* ∈ [*p*], we first c ompute t he p airwise E uclidean d istance m atrices f or t he *j* ^th^ and *k*^th^ genes,

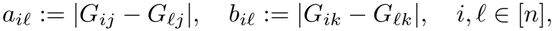

and then obtain the doubly-centered distances

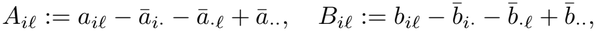

where *ā*_*i·*_ and *ā*_*·l*_ denote the *i*^th^ row mean and *l*^th^ column mean of (*a*_*il*_), respectively, and *ā*_*··*_ denotes the grand mean, with 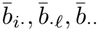 defined a nalogously f or (*b*_*il*_). The sample distance covariance is then given by

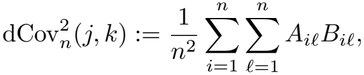

and the sample distance correlation coefficient is defined as

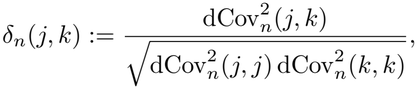

Provided 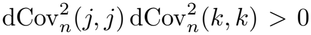, and *δ*_*n*_(*j, k*) := 0 otherwise. By construction, *δ*_*n*_(*j, k*) ∈ [0, 1], with values closer to 1 indicating stronger (possibly non-linear) dependence between *G*_*j*_ and *G*_*k*_. Distance correlation was computed with the *O*(*n* log *n*) algorithm of Huo and Székely [39] as implemented in the energy R package.

##### Mutual information

Mutual information provides a general, model-free measure of dependence that captures arbitrary (linear and non-linear) associations between two random variables. For every *j, k* ∈ [*p*], we consider the dependency measure

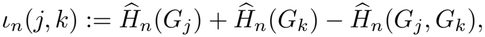

where *Ĥ*_*n*_(*G*_*j*_) and *Ĥ*_*n*_(*G*_*k*_) denote the estimated marginal entropies of the *j*^t h^ and *k*^th^ genes, respectively, and *Ĥ* (*G*, *G*) d enotes their e stimated joint e ntropy. Following the implementation in the minet R package [7], expression values are first discretized into 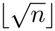 equal-frequency bins, and *Ĥ*_*n*_(*·*) is computed from the resulting frequency table using the empirical plug-in estimator. Similar to Chatterjee’s coefficient, *ι*_*n*_(*j, k*) is non-negative and equals zero if and only if *G*_*j*_ and *G*_*k*_ are independent, with larger values indicating stronger dependence. Unlike correlation coefficients, mutual information does not have an upper bound limit of 1.

For all measures, scores were converted to absolute values, symmetrized by taking the maximum of (*j, k*) and (*k, j*) where applicable, and set to 0 for undefined entries (e.g., constant genes).

#### 3.5.5 Data-driven calculation of testing cutoff

In the last two sections, we proposed two testing procedures, described in (7) and (8), for which, as previously indicated, it is necessary to determine the associated testing cutoffs in a data-driven way. Importantly, in our experiments, we observed that theoretical or asymptotic cutoffs, based on the impractical assumption of independence between the cellular gene expression observations, may lead to false results by either rejecting a significant proportion of true null hypotheses (anti-conservative or invalid tests) or not rejecting true alternatives (overly conservative and low-powered tests). To address this, we propose a general algorithm for estimating the test cutoff using a dataset, either existing or simulated, where the true null hypotheses are known a priori.

First, we fix any testing procedure to be established based on *n* cell observations, and for which the cutoff needs to be estimated. In this regard, we used a synthetic gene expression dataset, in which the number of cells is larger than *n*, and all true null hypotheses are known a priori, for example, the identity of all independent gene pairs in problem (1); gene pairs that are neither true edges nor true nulls (e.g., indirectly connected through a shared ancestor in the SERGIO simulations or an indirect path in STRING) are excluded. We denote the total number of cells by *n*′ ≥ *n*, and the total number of true nulls by *N*_0_. Now, we follow the steps formalized below and in Algorithm 1 to identify the ideal cutoff that controls the FPR.

1. Resample from the dataset a subsample of size *n*. In case *n* is reasonably smaller than *n*′, we resample without replacement; otherwise, especially when *n* = *n*′, we resample with replacement, as in the case of Bootstrap.
2. Fix some *C >* 0 as the testing cutoff and apply the testing procedure to the subsample. Moreover, let the total number of resulting false positives be denoted as *N*_fp_(*C*).
3. Record the following ratio:

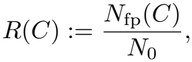

which clearly serves as an unbiased estimate of the FPR.
4. Since we desire to control the FPR below level *α*, find the maximum value as the testing cutoff *C* f or w hich *R* (*C*) i s k ept b elow *α*, a nd d enote i t a s t he optimal testing cutoff 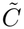, i.e.,

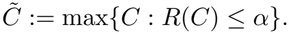

A grid search or some other suitable method is typically considered to find the above maxima.
5. Repeat the above steps 1-4 for *B* many iterations, and for every *b* ∈ [*B*], specifically denote the optimal testing cutoff, estimated in step 4, as 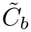.
6. Denote by *C*_*\**_ the median of all calculated optimal cutoffs 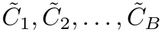, i.e.,

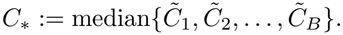

Consider *C*_*\**_ as the final testing cutoff, which clearly serves as an estimate of the true cutoff value. It is important to emphasize that the median is particularly chosen in the above to obtain a more robust estimate of the testing cutoff. More specifically, it prevents us from accidentally selecting a very large or small value of the final cutoff, which may o ccur as we estimate i t purely from the observational data. To control the FDR instead, we retain the scores of all labeled pairs (true edges and true nulls) from each of the *B* subsamples. For a cutoff *C*, we compute the false discovery proportion FDP(*C*) = *N*_fp_(*C*)*/* max {*N*_rej_(*C*), 1}, where *N*_rej_(*C*) is the number of pairs with score above *C*, and average it over the *B* subsamples. The FDR cutoff is the smallest *C* on a grid of 2000 score quantiles whose averaged FDP is at most *α* and whose average number of rejections is at least one, so that the largest admissible rejection set is retained. If no such cutoff exists, the FDR target is infeasible and no pair is called.

##### Algorithm 1

Cutoff estimation for testing dependence with sample size *n*

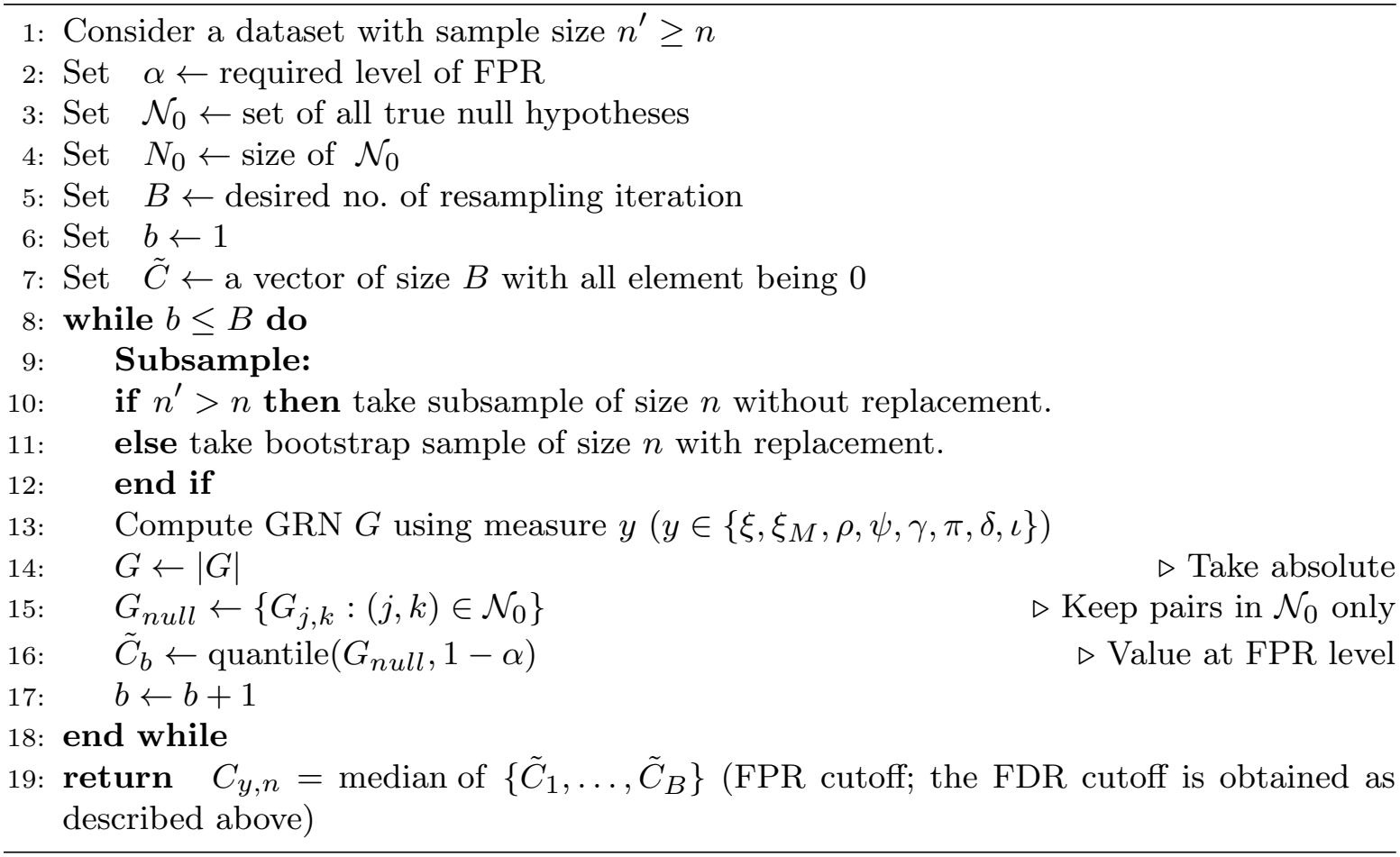

### 3.6 Testing for asymmetric predictive associations

In this section, we propose a multiple testing framework to test for *asymmetric predictive associations* between two dependent genes. To be specific, for every *j, k* ∈ [*p*], such that the *j*^th^ and *k*^th^ genes are dependent, *G*_*k*_ is said to be asymmetrically associated with *G*_*j*_ only if we can express *G*_*k*_ as *f*_*kj*_(*G*_*j*_) plus some possible, low-fluctuating noise term, for some nontrivial function *f*_*kj*_(*·*). In other words, *G*_*j*_ serves as an independent variable whereas *G*_*k*_ is dependent on *G*_*j*_ through some predictive function *f*_*kj*_(*·*). Moreover, if either of them shows asymmetric dependence on the other, we say that there is an asymmetric predictive association between these two genes. However, it is important to emphasize here that *G*_*j*_ and *G*_*k*_ can still be dependent without any asymmetric association between them, for example, when both are regulated simultaneously (co-regulated) by the expression of some third gene, or when they are linked by a monotone relationship, for which *ξ*_*n*_(*j* → *k*) and *ξ*_*n*_(*k* → *j*) are similar. As the coefficient measures statistical association, an asymmetric association indicates that one gene predicts the other, not that it causally regulates it.

We illustrate the notion of asymmetric (non-monotone) relationships in **Fig. 1a**, clearly distinguishing them from the cases of mere dependence.

#### 3.6.1 Multiple testing problem

Formally, for testing asymmetric predictive associations, we consider the following hypothesis testing problem: for every *j, k* ∈ [*p*], *j < k* such that the *G*_*j*_ and *G*_*k*_ are dependent,

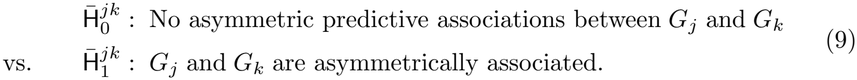

Here asymmetric predictive association means either the expression of *G*_*j*_ depends on *G*_*k*_, or vice versa. This extends the problem from just testing for dependence to testing for asymmetric predictive associations among two genes, which, to the best of our knowledge, has not been considered in past literature.

#### 3.6.2 Testing procedure

To propose a solution to the testing problem in (9), first, it is important to consider a dependency measure that can suitably capture the notion of asymmetric predictive associations. More specifically, for every *j, k* ∈ [*p*] such that *G*_*j*_ and *G*_*k*_ are dependent, we must measure the extent to which *G*_*k*_ is predicted by *G*_*j*_, i.e., to what degree *G*_*k*_ can be expressed in terms of *G*_*j*_ through some function. The definition of Chatterjee’s correlation coefficient *ξ*_*n*_(*j* → *k*), as defined in (2), or (3), makes it suitable for testing for asymmetric associations. In fact, following [11, Theorem 1.1] Chatterjee’s coefficient converges to 1 if and only if *G*_*k*_ = *f*_*kj*_(*G*_*j*_) for some measurable function *f*_*kj*_(*·*), i.e., *G*_*k*_ is completely dependent on *G*_*j*_. Therefore, if *ξ*_*n*_(*j* → *k*) is significantly larger than *ξ*_*n*_(*k* → *j*), it serves as strong evidence that *G*_*k*_ is asymmetrically associated with, and predicted by, *G*_*j*_ whereas the degree of the opposite prediction is comparatively weak. Meanwhile, if both *ξ*_*n*_(*k* → *j*) and *ξ*_*n*_(*j* → *k*) are simultaneously large, i.e., similar to each other, that suggests that both genes are related monotonically (e.g., linearly) or are co-regulated by some third gene contributing to higher values in both directions.

Thus, based on the above discussion and following the rationales stated in Section 3.5.3, we propose the following testing procedure,

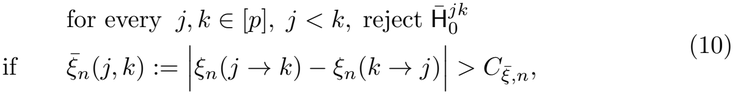

where 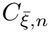 is some positive constant, which must be appropriately determined such that it controls the FPR at the required level. However, since any characterization of the limiting behavior of (*ξ*_*n*_(*j* → *k*) − *ξ*_*n*_(*k* → *j*)) is not present in the theoretical literature even for IID observations, we need to estimate 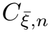 from simulated or curated gene expression datasets, where the monotone (null) and non-monotone gene pairs are a priori specified. The same test is also applied with ChatterjeeKNN, replacing *ξ*_*n*_ by *ξ*_*n*,*M*_ in (10).

It is noteworthy that GRNBoost2 generates an asymmetric dependency measure provided by the elements of the asymmetric importance matrix *W* evaluated at the tree construction step. Therefore, in a similar vein, we propose the following testing procedure,

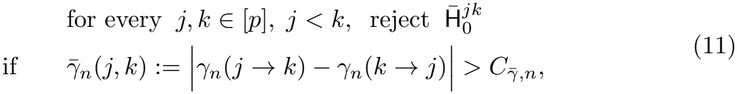

which we later compare with (10) in Section 2.2. Now, we follow the steps formalized below in Algorithm 2 to identify the ideal cutoff for the testing problems (10) and (11) that controls the FPR at a defined level.

##### Algorithm 2

Cutoff estimation for testing asymmetric predictive associations on sample size *n*

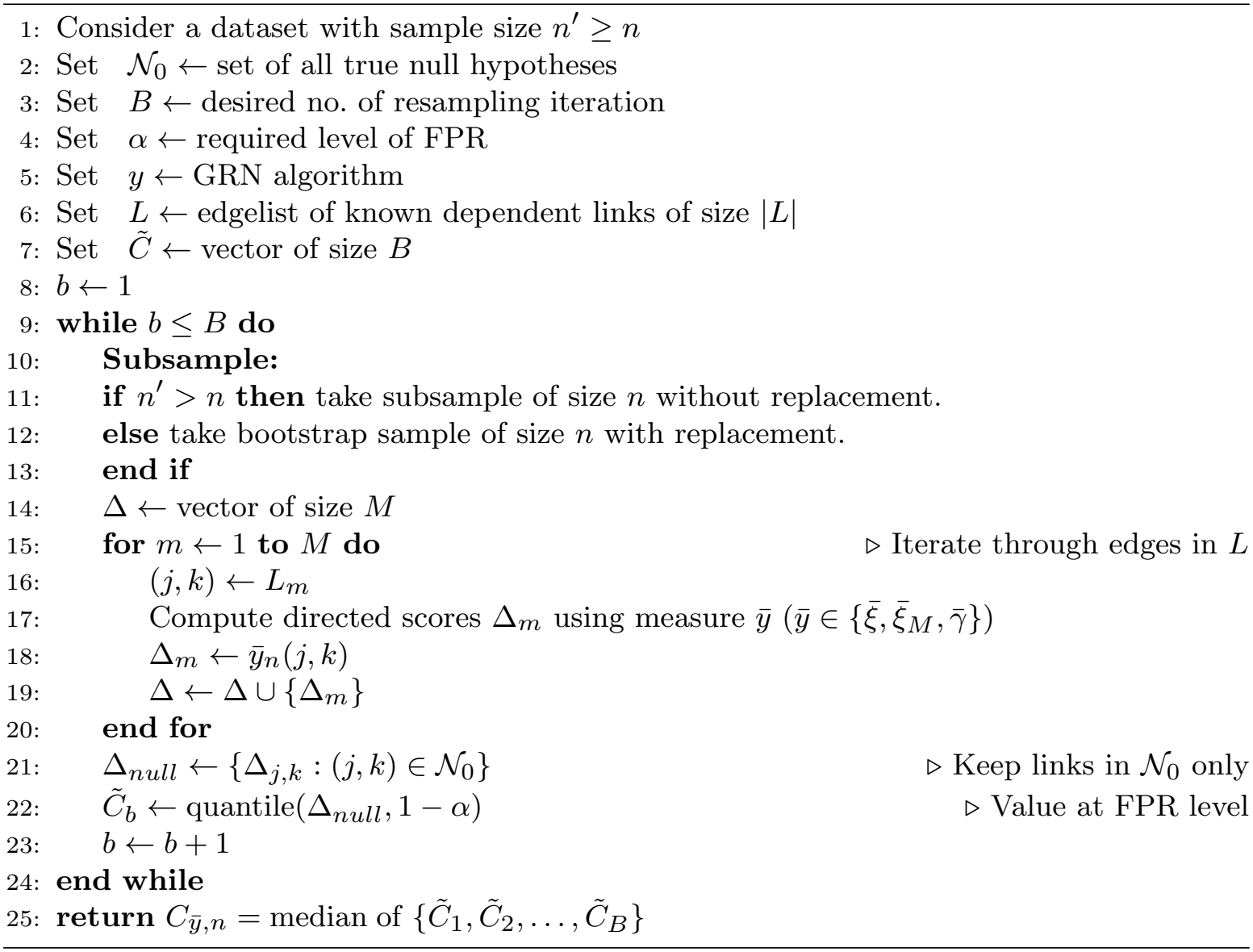

In the simulations, the null set *N*_0_ consists of the monotone (linear and exponential) pairs. To control the FDR instead, we rank the labeled pairs of each subsample by Δ_*m*_ and take the largest rejection set whose false discovery proportion is at most *α*; the cutoff is the midpoint between the last rejected and first non-rejected score, and the final FDR cutoff is the median over the *B* subsamples. If no non-empty set qualifies, the maximum score is used, so that nothing is rejected.

On real data, *N*_0_ consists of dependent gene pairs that are subunits of the same CORUM complex and have no TFLink annotation in either direction. Pairs are the dependent pairs found in the target dataset, and the cutoff is learned directly on that dataset using *n* = 1500 cells subsampled without replacement from all its cells, *B* = 50, and the FPR criterion.

### 3.7 Stability and transferability of the learned cutoff

#### 3.7.1 Stability with sample size

To test whether the cutoff learned by Algorithm 2 is stable as the amount of data grows, we used the generative model of Section 3.4 (500 genes, 250 gene pairs, noise standard deviation 0.5, dropout rate 0.3, mean expression 15) with the total number of cells varying from 500 to 10,000 in steps of 500, and 10 replicates per size. For each replicate, 60% of the cells formed the training set and 40% the test set, so the test size *n* ranged from 200 to 4,000. The cutoff was learned on the training set with *n*′ equal to the training size, subsampling *n* cells without replacement, *B* = 50 and *α* = 0.05, once with the FPR criterion and once with the FDR criterion, with the monotone pairs as the null set *N*_0_. The learned cutoffs were then applied unchanged to the held-out test cells, and we recorded the realized FPR, FDR and sensitivity. To summarize the trend of the cutoff with *n*, we fitted a line to the median cutoff per *n* on the log-log scale; this fit is descriptive and is not used to set cutoffs in any other analysis, except as the baseline comparator in **Supplementary Fig. 7a**.

#### 3.7.2 Transfer across data characteristics

To test whether a cutoff can be transferred between datasets, we simulated data in which one characteristic was changed at a time from a baseline setting (noise standard deviation 0.5, dropout rate 0.3, mean expression 15, gene-mean dispersion 0.5, Poisson counts; 500 genes, 250 pairs). The settings varied were the noise standard deviation (0.2, 1, 1.5), the dropout rate (0, 0.1, 0.2, 0.5), the mean expression (10, 20, 30), the dispersion (0.2, 1, 1.5), and the count distribution (negative binomial with size 2 instead of Poisson). The gene pairs were identical across settings, and the interaction noise was drawn independently for each replicate. For each setting, we generated 5 replicates at 1,000, 2,000, 3,750, 5,000 and 7,500 cells. One replicate with 7,500 cells served as the reference (*n*′ = 7500). In each setting, Algorithm 2 (FPR criterion, *B* = 50, *α* = 0.05, monotone pairs as *N*_0_) was run on the reference to learn a cutoff for *n* = 1000, 2000 and 3750. The cutoff was then applied unchanged to four other replicates of the same setting at that *n*, and the realized FPR was recorded. For cross-setting transfer, the cutoff learned in one setting was applied to the held-out replicates of every other setting at *n* = 2000, and the realized FPR was averaged over replicates.

Both analyses used ChatterjeeKNN (*M* = 5) with the asymmetry score |*ξ*_*n*,*M*_ (*j* → *k*) − *ξ*_*n*,*M*_ (*k* → *j*)| of (10).

### 3.8 Testing to identify dependency on real data

For the proposed procedures in (7) and (8), which consider testing of independence through the problem in (1), we consider the human protein-protein interaction (PPI) network from the STRING database [24, 25], a comprehensive collection of text-mined and experimentally determined functional associations, as an estimated ground truth. We kept only interactions with a combined score greater than 900. Gene pairs were labeled using the high-confidence graph over the full proteome: pairs directly connected were taken as true edges; pairs with no path shorter than three steps (including genes with no high-confidence interaction) were taken as true nulls; and pairs two steps apart were excluded from calibration and evaluation, as they are likely indirectly dependent. Since the cutoff for Chatterjee’s correlation depends on *n*, we estimated a single testing cutoff from one reference dataset (GSE239592) using Algorithm 1 over *B* = 50 iterations, and applied it without modification to the remaining target datasets. At each iteration, we drew a subsample of 1500 cells without replacement from all cells of the reference dataset to compute the empirical cutoff at the desired FPR level of 0.05. The median cutoff across iterations was taken as the final value. Cells were maintained at 1500 throughout, for both the reference dataset used to estimate the cutoff and each target dataset to which the cutoff was subsequently applied (a fixed random subsample of 1500 cells per dataset). We used the common gene panel of 674 genes detected in more than 30% of T cells in every dataset (Section 3.1), and computed ChatterjeeKNN (*M* = 5) on the log-normalized expression values. The cutoff estimated from the reference dataset was then applied to each target dataset to identify significantly dependent gene pairs. Realized FPR and FDR in each dataset were computed over the labeled pairs.

### 3.9 Testing to identify asymmetric dependence on real data

Since, to the best of our knowledge, there is no database in real-world settings with the ground truth of asymmetric associations clearly specified, we learned the cutoff for the test (10) directly on the target dataset (GSM8331607) with Algorithm 2, using a curated proxy for the null set. The known links *L* were the gene pairs found dependent in the target dataset by the dependency test of the previous subsection. The null set *N*_0_ consisted of those dependent pairs whose genes are subunits of the same protein complex in CORUM [26], excluding any pair with a TFLink annotation in either direction [27]. Subunits of a stable complex are co-regulated and expected to be related symmetrically, so neither gene should predict the other asymmetrically. The cutoff was learned with *n* = 1500 cells subsampled without replacement from all cells of the target dataset (*n*′ *> n*), *B* = 50, *α* = 0.05, and the FPR criterion, and applied unchanged to the same 1500-cell subsample. The asymmetry score was Δ_*jk*_ = *ξ*_*n*,*M*_ (*j* → *k*) − *ξ*_*n*,*M*_ (*k* → *j*) with ChatterjeeKNN (*M* = 5), and a pair was called asymmetrically associated if |Δ_*jk*_| *> C*. For called pairs, the predictor was the gene with the larger *ξ*_*n*,*M*_ (*·* → *·*), i.e., *j* if Δ_*jk*_ ≥ 0. We checked the orientation of the implementation on simulated data (*y* = *x*^2^) before use.

To assess the inferred direction, we used the transcription factor (TF)–target pairs in TFLink (human, all evidence types, v1.0). Among called pairs annotated in TFLink in either direction, a call was counted as correct if the predictor–target direction matched the annotated TF-to-target direction. Ranked expression scatter plots (with LOESS smoothing) were generated for the top-ranked pair by |Δ_*jk*_| and for the genes in the ten strongest asymmetric pairs.

### 3.10 Performance metrics

For any testing procedure, we evaluate its testing and learning performances in terms of some standard performance metrics. Specifically, we evaluate their testing performance by using the following performance metrics:

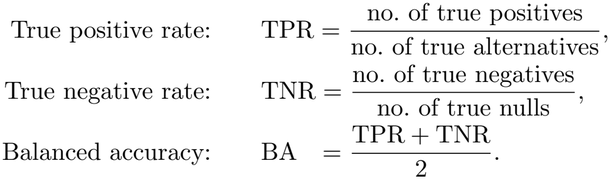

Besides, we evaluate the performance of their classification by using AUROC, the area under the ROC curve that plots TPR vs FPR.We also determined the learning performance of [0, 1]-bounded algorithms by computing the root mean square error (RMSE).

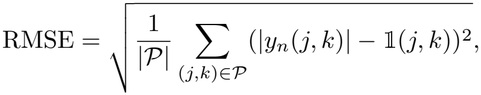

where *y* ∈ {*ξ, ξ*_*M*_, *ρ, ψ, δ*}, |*y*_*n*_(*j, k*)| is the absolute symmetrized score, which already lies in [0, 1] for these measures, and *P* is the set of labeled gene pairs (*j, k*), *j < k*, i.e., true edges and true nulls, excluding pairs removed from evaluation (e.g., trek pairs in the SERGIO analysis of Supplementary Fig. 5 or two-step STRING pairs). RMSE was not computed for pcNet, GRNBoost2 or mutual information, whose raw scores are unbounded and not directly comparable to the ground-truth indicator on this scale. Here, 1 (*j, k*) denotes the indicator function that takes value 1 if and only if (*j, k*) is a true edge in the ground truth, i.e., 1 (*j, k*) := 1 *{*(*j, k*) is a true edge*}*.

## 4 Discussion

In this study, we introduced a multiple-testing framework based on Chatterjee’s rank correlation coefficient for constructing GRNs from scRNA-seq data, and demonstrated its effectiveness across simulated and real datasets. Through a series of simulation studies, we first established that Chatterjee’s correlation is more effective at capturing gene-gene dependencies than the classical correlation coefficients (Pearson, Spearman). Notably, it also compared favorably against two established nonparametric measures, distance correlation and mutual information, that, like Chatterjee’s coefficient, are capable of detecting non-linear associations without distributional assumptions. This places our method within a broader class of rank- and information-based dependency measures increasingly used in transcriptomic network inference. Within this class, Chatterjee’s coefficient achieved competitive or superior sensitivity and balanced accuracy at substantially lower computational cost, since it operates on a single, closed-form rank statistic rather than requiring density or distance-matrix estimation.

We then used a more complex simulation framework, SERGIO, to model intricate, trajectory-driven gene interactions. In this more challenging setting, Chatterjee’s correlation matched or surpassed the traditional and nonparametric correlation measures under FPR control and showed promising behavior relative to more advanced machine-learning algorithms, including regression tree-based GRNBoost2 [8, 40] and manifold-alignment-based PCNet [9]. We also evaluated ChatterjeeKNN, the nearest-neighbor extension of the coefficient [12], whose advantage was greatest for directional inference. In the asymmetric association test, it detected substantially more true asymmetric pairs than both the standard coefficient and GRNBoost2. Its sensitivity was unchanged under FDR control, whereas that of GRNBoost2 halved. Among detected pairs, it recovered the correct direction almost without exception, matching GRN-Boost2 and exceeding the standard coefficient, with less variability across replicates. It also achieved the lowest RMSE among the bounded measures in every simulation. The standard coefficient retained higher sensitivity for dependence detection under FPR-controlled cutoffs in some SERGIO datasets, and ChatterjeeKNN was slower than the standard coefficient while remaining faster than distance correlation. Given its gains in directional inference, we used ChatterjeeKNN for the real-data analyses; its parameter *M* was chosen on simulated data, and performance was insensitive to *M* for *M* ≤ 10.

We deliberately benchmarked Chatterjee’s correlation against a broad spectrum of tools spanning classical parametric correlation, nonparametric symmetric dependency measures, and machine-learning-based network reconstruction methods, in order to situate it comprehensively within the existing GRN inference landscape. We recognize that this comparison across categories is not entirely like-for-like: GRNBoost2 and PCNet leverage global, multivariate information across the full gene expression matrix, whereas Chatterjee’s correlation, like the other pairwise measures considered here, evaluates each gene pair independently and does not model higher-order network structure. Despite this difference in scope, we considered it necessary to include these machine-learning methods as comparators, since they represent the current de facto standard for GRN reconstruction from snapshot, non-perturbed, single-cell data and provide essential context for evaluating whether a simple, transparent, pairwise statistic can remain competitive with more complex, global approaches. Our results show that Chatterjee’s correlation remains competitive, although not uniformly, as GRNBoost2 outperformed Chatterjee’s correlation in sensitivity, balanced accuracy, and AUROC in the custom simulation. Chatterjee’s correlation nevertheless retains the interpretability and computational simplicity of a single pairwise rank statistic.

A key advantage of our approach is its ability to detect asymmetric (directional) dependence within GRNs. By exploiting the non-symmetric form of Chatterjee’s correlation, we were able to identify predictor → response relationships directly from a pairwise statistic, without requiring a global model of the network. We tested this by simulating GRNs with both non-monotone (parabolic, sinusoidal) and monotone (linear, exponential) interactions, and our method proved more sensitive than GRN-Boost2 in detecting asymmetric pairs. This directionality, however, is intrinsically tied to non-monotonicity: Chatterjee’s coefficient is asymmetric only when the underlying dependency is non-monotone (many-to-one), and this asymmetry vanishes under monotone relationships, whether linear or non-linear. Consequently, our approach cannot resolve the direction of dependence between genes whose expression is related in a monotone fashion. This is a substantive limitation, because many regulatory responses, such as dose-dependent activation or repression, are expected to be monotone. Still, directionality calls only apply to the non-monotone part of a gene pair’s dependency, not to the association as a whole.

Our proposed cutoff selection procedure, while effective, is currently reliant on a well-curated ground-truth network. In this study, we used the STRING database to provide this ground truth for the dependency test and protein complexes from CORUM as the null set for the asymmetric association test in our real-data application. We acknowledge that the performance of this approach is tied to the quality of the reference network, and that transferring a cutoff estimated from one dataset to another assumes a degree of similarity in cell-cell dependency structure between the two. Our simulations confirmed this: learned cutoffs were stable with sample size but did not transfer between datasets that differed in dropout, noise, or dispersion, so the cutoff should be relearned when data characteristics differ, and we checked that the four PBMC datasets had broadly similar distributions before transferring the cut-off. A related limitation concerns FDR control. In simulations with complete ground truth, FDR-controlled cutoffs could be estimated, although in SERGIO this required excluding indirectly dependent (trek) gene pairs from the null set. On real data, FDR could not be controlled, because high-confidence STRING interactions are sparse and incomplete, so genuine dependencies without a recorded interaction are counted as false positives. We therefore calibrated the real-data tests to control the FPR, which depends only on the null distribution. However, the STRING database is continuously updated, reflecting the latest advances in biological knowledge. We believe that as such databases become more comprehensive and accurate, our method will become even more effective and scalable, offering a more robust way to identify significant regulatory interactions in gene networks.

More fundamentally, asymmetric dependence does not establish regulatory or causal direction. A larger *ξ*_*n*_(*j* → *k*) than *ξ*_*n*_(*k* → *j*) indicates that *G*_*k*_ is more nearly a function of *G*_*j*_ than vice versa, which is compatible with *G*_*j*_ regulating *G*_*k*_ but can also arise from non-invertible relationships, unequal noise or marginal distributions, latent common causes, mixtures of cell states, selection, and dropout or other measurement error. For example, the U-shaped LTB–S100A4 relationship in our PBMC analysis is also consistent with a mixture of CD4^+^ and CD8^+^ T cells. The directed edges reported here should therefore be read as hypotheses that require experimental validation, such as perturbation studies.

An alternative approach to inferring directed gene regulation is via Bayesian causal structure learning over DAGs [18–22], which offers a principled, model-based frame-work for global causal discovery but at substantial computational cost and with identifiability challenges that grow super-exponentially with the number of genes. To the best of our knowledge, our framework is the only method for directed GRN inference from scRNA-seq data that is simultaneously nonparametric and non-Bayesian: it requires no distributional assumptions on gene expression, no prior specification, and no global network model, relying instead on a simple, closed-form, pairwise rank statistic. This positions our approach as a lightweight, model-free complement to Bayesian DAG learning, well suited either as a standalone tool for directionally informed GRN construction or as a fast pre-screening step that narrows the search space for more computationally intensive causal discovery methods.

Taken together, our findings underscore the utility of Chatterjee’s correlation for accurately identifying and directing asymmetric predictive associations in complex biological networks that exhibit non-monotone dependencies, while remaining computationally efficient, distribution-free, and interpretable, properties that are increasingly important as single-cell datasets continue to grow in scale and complexity.

## 5 Declarations

### 5.3 Data Availability

The sources of the data sets used in this article can be found in **Supplementary Table 1**. Simulated datasets using SERGIO were obtained from https://github.com/PayamDiba/SERGIO. The ground truth PPI network was obtained from the STRING database v12.0 (https://string-db.org/), the transcription factor–target annotations were obtained from the TFLink database v1.0 (human, all evidence types; https://tflink.net/), and the human protein complexes used as the null set for the asymmetric association test were obtained from the CORUM database (https://mips.helmholtz-muenchen.de/corum/).

### 5.4 Code Availability

All code used to generate the results in this study is available at https://github.com/Xenon8778/scXiNet.

### 5.5 Author contributions

S.G.: Conceptualization, Methodology, Visualization, Software, Formal Analysis, Writing - Original Draft. A.C.: Conceptualization, Methodology, Formal Analysis, Writing - Original Draft. V.R.: Software, Formal Analysis. Y.N.: Writing - Review & Editing. J.J.C: Conceptualization, Supervision, Writing - Review & Editing, Resources.

## References

[1] Blinov, M.L., Faeder, J.R., Goldstein, B., Hlavacek, W.S.: A network model of early events in epidermal growth factor receptor signaling that accounts for combinatorial complexity. Biosystems 83(2-3), 136–151 (2006)

[2] Waters, K.M., Liu, T., Quesenberry, R.D., Willse, A.R., Bandyopadhyay, S., Kathmann, L.E., Weber, T.J., Smith, R.D., Wiley, H.S., Thrall, B.D.: Network analysis of epidermal growth factor signaling using integrated genomic, proteomic and phosphorylation data. PloS one 7(3), 34515 (2012)

[3] Blencowe, M., Arneson, D., Ding, J., Chen, Y.-W., Saleem, Z., Yang, X.: Network modeling of single-cell omics data: challenges, opportunities, and progresses. Emerging topics in life sciences 3(4), 379–398 (2019)

[4] Haque, A., Engel, J., Teichmann, S.A., Lönnberg, T.: A practical guide to single-cell rna-sequencing for biomedical research and clinical applications. Genome medicine 9(1), 75 (2017)

[5] De Smet, R., Marchal, K.: Advantages and limitations of current network inference methods. Nature Reviews Microbiology 8(10), 717–729 (2010)

[6] Székely, G.J., Rizzo, M.L., Bakirov, N.K.: Measuring and testing dependence by correlation of distances. Ann. Statist. 35(1), 2769–2794 (2007)

[7] Meyer, P.E., Lafitte, F., Bontempi, G.: minet: Ar/bioconductor package for inferring large transcriptional networks using mutual information. BMC bioinformatics 9(1), 461 (2008)

[8] Moerman, T., Aibar Santos, S., Bravo González-Blas, C., Simm, J., Moreau, Y., Aerts, J., Aerts, S.: Grnboost2 and arboreto: efficient and scalable inference of gene regulatory networks. Bioinformatics 35(12), 2159–2161 (2019)

[9] Osorio, D., Zhong, Y., Li, G., Huang, J.Z., Cai, J.J.: sctenifoldnet: a machine learning workflow for constructing and comparing transcriptome-wide gene regulatory networks from single-cell data. Patterns 1(9) (2020)

[10] Kang, Y., Thieffry, D., Cantini, L.: Evaluating the reproducibility of single-cell gene regulatory network inference algorithms. Frontiers in genetics 12, 617282 (2021)

[11] Chatterjee, S.: A new coefficient of correlation. Journal of the American Statistical Association 116(536), 2009–2022 (2021)

[12] Lin, Z., Han, F.: On boosting the power of chatterjee’s rank correlation. Biometrika 110(2), 283–299 (2023)

[13] Sadeghi, B.: Chatterjee correlation coefficient: A robust alternative for classic correlation methods in geochemical studies-(including “triplecpy” python package). Ore Geology Reviews 146, 104954 (2022)

[14] Suo, C., Polanski, K., Dann, E., Lindeboom, R.G., Vilarrasa-Blasi, R., Vento-Tormo, R., Haniffa, M., Meyer, K.B., Dratva, L.M., Tuong, Z.K., et al.: Dandelion uses the single-cell adaptive immune receptor repertoire to explore lymphocyte developmental origins. Nature Biotechnology 42(1), 40–51 (2024)

[15] Glymour, C., Zhang, K., Spirtes, P.: Review of causal discovery methods based on graphical models. Frontiers in genetics 10, 524 (2019)

[16] Kelly, J., Berzuini, C., Keavney, B., Tomaszewski, M., Guo, H.: A review of causal discovery methods for molecular network analysis. Molecular genetics & genomic medicine 10(10), 2055 (2022)

[17] Chaudhuri, A., Bhattacharya, A., Ni, Y.: MARCEDES: Score-based causal discovery under non-gaussianity with continuous optimization. arXiv preprint arXiv:2609.30643 (2026)

[18] Choi, J., Chapkin, R., Ni, Y.: Bayesian causal structural learning with zero-inflated Poisson Bayesian networks. Advances in Neural Information Processing Systems 33, 5887–5897 (2020)

[19] Choi, J., Ni, Y.: Model-based causal discovery for zero-inflated count data. Journal of Machine Learning Research 24(200), 1–32 (2023)

[20] Choi, J., Chapkin, R.S., Ni, Y.: Bayesian differential causal directed acyclic graphs for observational zero-inflated counts with an application to two-sample single-cell data. The Annals of Applied Statistics 19(3), 1908 (2025)

[21] Chaudhuri, A., Bhattacharya, A., Ni, Y.: Consistent DAG selection for bayesian causal discovery under general error distributions. arXiv preprint arXiv:2508.00993 (2025)

[22] Chaudhuri, A., Ni, Y., Bhattacharya, A.: Consistent bayesian causal discovery for structural equation models with equal error variances. arXiv preprint arXiv:2509.15197 (2025)

[23] Dibaeinia, P., Sinha, S.: Sergio: a single-cell expression simulator guided by gene regulatory networks. Cell systems 11(3), 252–271 (2020)

[24] Mering, C.v., Huynen, M., Jaeggi, D., Schmidt, S., Bork, P., Snel, B.: String: a database of predicted functional associations between proteins. Nucleic acids research 31(1), 258–261 (2003)

[25] Szklarczyk, D., Kirsch, R., Koutrouli, M., Nastou, K., Mehryary, F., Hachilif, R., Gable, A.L., Fang, T., Doncheva, N.T., Pyysalo, S., et al.: The string database in 2023: protein–protein association networks and functional enrichment analyses for any sequenced genome of interest. Nucleic acids research 51(D1), 638–646 (2023)

[26] Giurgiu, M., Reinhard, J., Brauner, B., Dunger-Kaltenbach, I., Fobo, G., Frishman, G., Montrone, C., Ruepp, A.: Corum: the comprehensive resource of mammalian protein complexes—2019. Nucleic acids research 47(D1), 559–563 (2019)

[27] Liska, O., Bohár, B., Hidas, A., Korcsmáros, T., Papp, B., Fazekas, D., Ari, E.: Tflink: an integrated gateway to access transcription factor–target gene interactions for multiple species. Database 2022, 083 (2022)

[28] Müller, J.R., Siebenlist, U.: Lymphotoxin beta receptor induces sequential activation of distinct nf-κb factors via separate signaling pathways. Journal of Biological Chemistry 278(14), 12006–12012 (2003)

[29] Wang, Y.-f., Zhang, W.-l., Li, Z.-x., Liu, Y., Tan, J., Yin, H.-z., Zhang, Z.-c., Piao, X.-j., Ruan, M.-h., Dai, Z.-h., et al.: Mettl14 downregulation drives s100a4+ monocyte-derived macrophages via myd88/nf-κb pathway to promote mafld progression. Signal transduction and targeted therapy 9(1), 91 (2024)

[30] Wang, X., Ni, Y., Shi, Y., Yu, X., Wu, C., Xiang, Z., Lv, W., Wang, Q.: Exploring ltb-mediated t cell differentiation as a prognostic marker in triple-negative breast cancer. Breast Cancer Research: BCR (2026)

[31] Weatherly, K., Bettonville, M., Torres, D., Kohler, A., Goriely, S., Braun, M.Y.: Functional profile of s100a4-deficient t cells. Immunity, inflammation and disease 3(4), 431–444 (2015)

[32] Hayashi, F., Mittl, K., Dandekar, R., Gerdts, J., Hassan, E., Schubert, R.D., Oshiro, L., Loudermilk, R., Greenfield, A., Augusto, D.G., et al.: Antigen specificity of clonally enriched cd8+ t cells in multiple sclerosis. Nature Immunology 27(3), 490–502 (2026)

[33] Hao, Y., Stuart, T., Kowalski, M.H., Choudhary, S., Hoffman, P., Hartman, A., Srivastava, A., Molla, G., Madad, S., Fernandez-Granda, C., et al.: Dictionary learning for integrative, multimodal and scalable single-cell analysis. Nature biotechnology 42(2), 293–304 (2024)

[34] Germain, P.-L., Lun, A., Meixide, C.G., Macnair, W., Robinson, M.D.: Doublet identification in single-cell sequencing data using scdblfinder. f1000research 10, 979 (2022)

[35] Sun, W., Tony Cai, T.: Large-scale multiple testing under dependence. Journal of the Royal Statistical Society Series B: Statistical Methodology 71(2), 393–424 (2009)

[36] Leek, J.T., Storey, J.D.: A general framework for multiple testing dependence. Proceedings of the National Academy of Sciences 105(48), 18718–18723 (2008)

[37] Chaudhuri, A., Fellouris, G.: Sequential detection and isolation of a correlated pair. In: 2020 IEEE International Symposium on Information Theory (ISIT), pp. 1141–1146 (2020). IEEE

[38] Chaudhuri, A., Fellouris, G.: Joint sequential detection and isolation for dependent data streams. The Annals of Statistics 52(5), 1899–1926 (2024)

[39] Huo, X., Székely, G.J.: Fast computing for distance covariance. Technometrics 58(4), 435–447 (2016)

[40] Breiman, L.: Random forests. Machine learning 45(1), 5–32 (2001)

