## Supplementary Figures for "Constructing Gene Regulatory Network using Chatterjee’s Rank Correlation with Single-cell Transcriptomic Data"

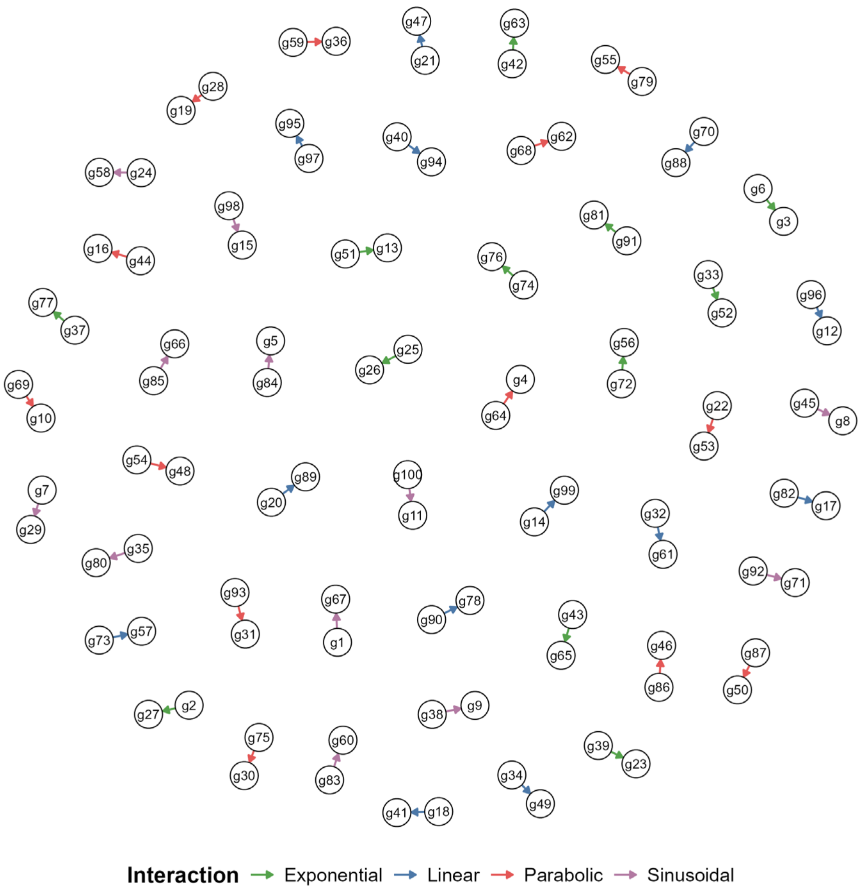

**Supplementary Figure 1.** Ground-truth network topology of the simulated scRNA-seq dataset.

The network comprises 50 disjoint gene pairs, each linked by a directed edge from a source gene to a target gene. Edge colors denote the interaction type used to generate the target gene's expression: exponential, linear, parabolic, or sinusoidal.

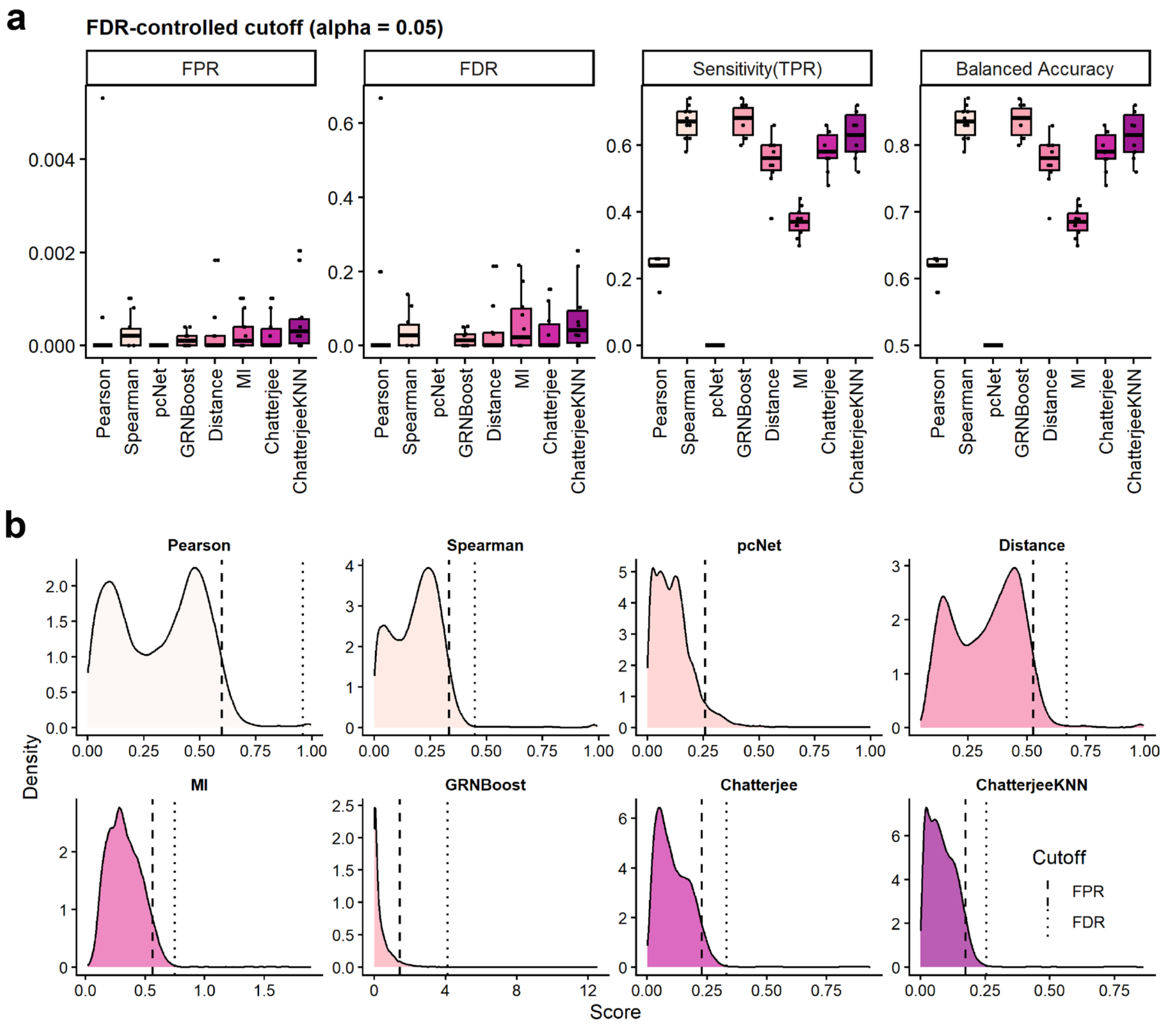

**Supplementary Figure 2.** Performance benchmarking and score distributions under FDR-controlled cutoffs.

(**a**) Performance of eight GRN inference methods across test datasets using the following classification metrics: FPR, FDR, sensitivity (TPR), and balanced accuracy, with cutoffs controlling the false discovery rate (FDR) at $\alpha = 0.05$. (**b**) Density distributions of inferred links from test data and their respective optimal cutoffs. Thresholds were computed from the training data to control the FPR (dashed line) or the FDR (dotted line) at 5% across our simulated scRNA-seq datasets featuring non-linear dependencies.

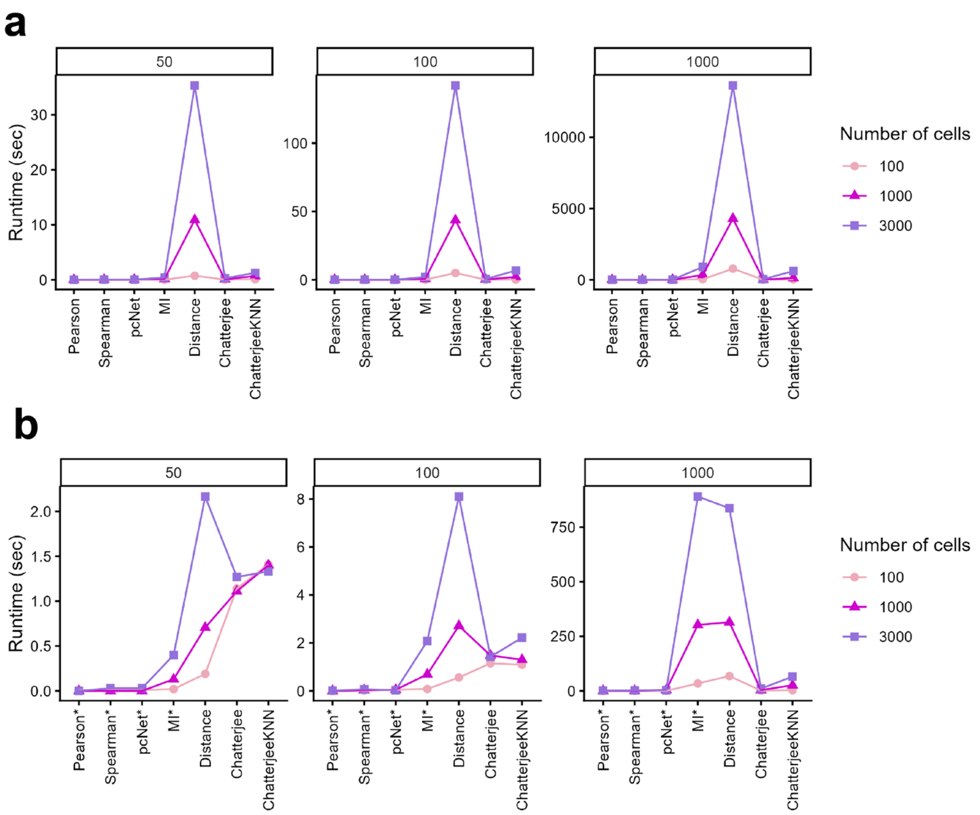

**Supplementary Figure 3.** Empirical runtimes of GRN inference algorithms across varying dataset sizes.

Computational execution times were evaluated using simulated scRNA-seq datasets across combinations of different gene counts (50, 100, 1000) and cell numbers (100, 1000, 3000). (a) Runtimes with all methods run on a single core. (b) Runtimes with Distance correlation, Chatterjee, and ChatterjeeKNN parallelized across 8 cores; the other methods are unchanged.

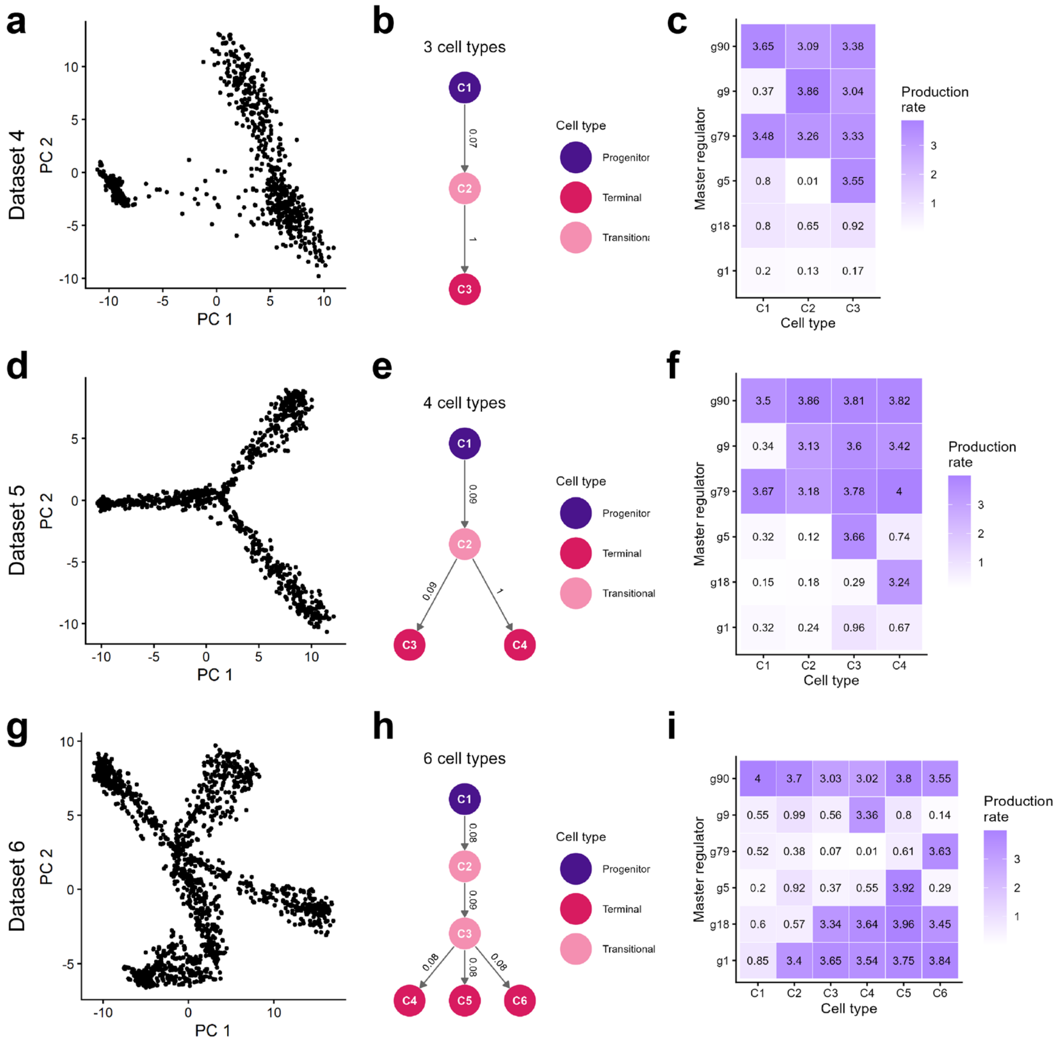

**Supplementary Figure 4.** Differentiation structure of the SERGIO datasets DS4-DS6.

Rows correspond to datasets, and columns show the PCA, differentiation graph, and master regulator production rates. (**a-c**) DS4: (**a**) PCA of the simulated cells showing a linear trajectory, (**b**) differentiation graph with 3 cell types, and (**c**) production rates of the six master regulators (g90, g9, g79, g5, g18, g1) in each cell type. (**d-f**) DS5: (**d**) PCA showing a bifurcating trajectory, (**e**) differentiation graph with 4 cell types, and (**f**) master regulator production rates. (**g-i**) DS6: (**g**) PCA showing a trifurcating trajectory, (**h**) differentiation graph with 6 cell types, and (**i**) master regulator production rates. In the differentiation graphs, nodes are cell types colored by role (progenitor, transitional, or terminal) and edge labels give cell-type migration rates.

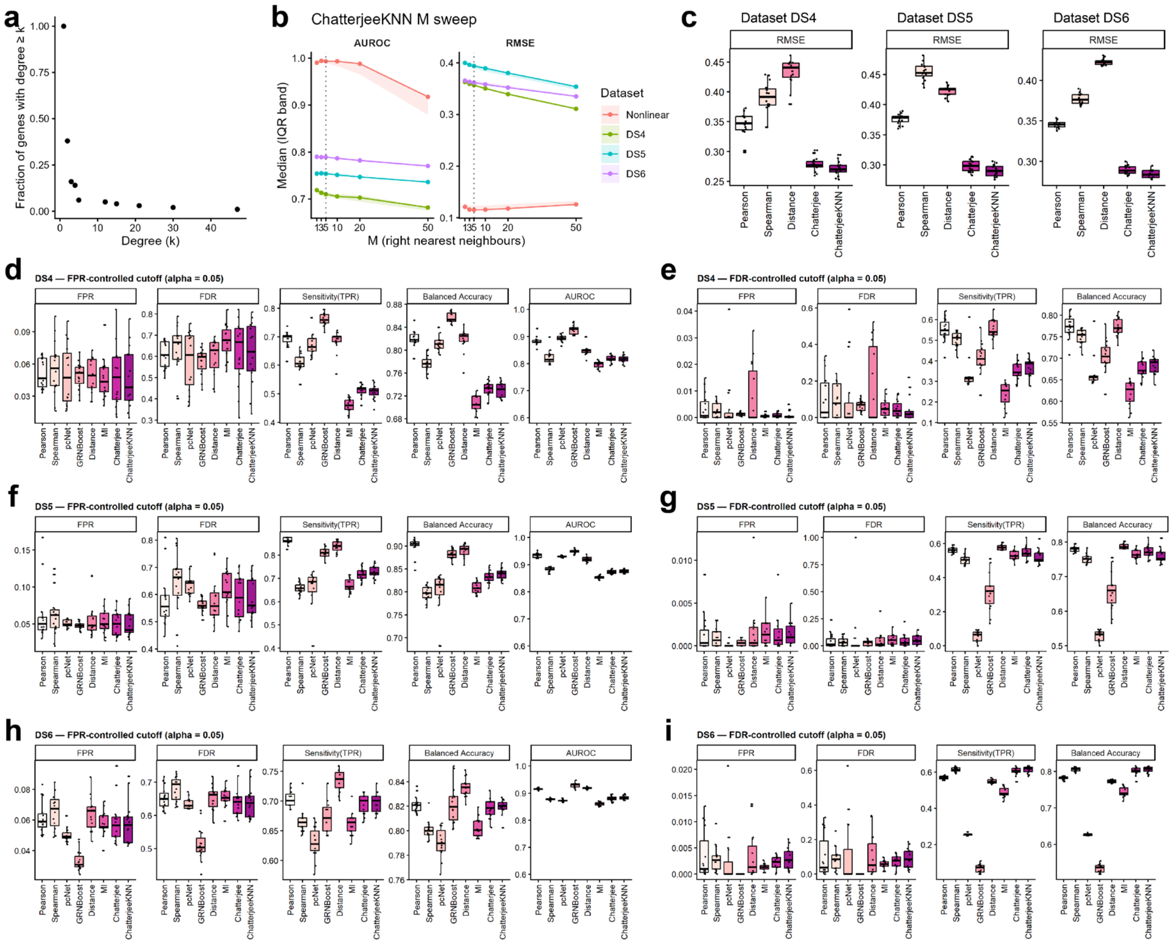

**Supplementary Figure 5.** Benchmarking on SERGIO datasets with trek pairs excluded from the null set, and selection of $M$ for ChatterjeeKNN.

(a) Degree distribution of the DS4-DS6 ground-truth GRN, shown as the fraction of genes with degree $\geq k$. (b) Selection of $M$, the number of right nearest neighbors in ChatterjeeKNN. AUROC and RMSE are shown as a function of $M$ for the non-linear simulation and for DS4, DS5, and DS6. Lines show the median and bands the interquartile range over five replicates, and the dotted line marks the chosen value, $M = 5$. (c) RMSE of the [0,1] bounded dependency metrics (Pearson, Spearman, Distance, Chatterjee, and ChatterjeeKNN correlation) for DS4, DS5, and DS6. (d, f, h) Classification performance (FPR, FDR, sensitivity (TPR), balanced accuracy, and AUROC) of eight network inference methods with FPR-controlled cutoffs at $\alpha = 0.05$ for DS4 (d), DS5 (f), and DS6 (h). (e, g, i) The same metrics, except AUROC, with FDR-controlled cutoffs at $\alpha = 0.05$ for DS4 (e), DS5 (g), and DS6 (i).

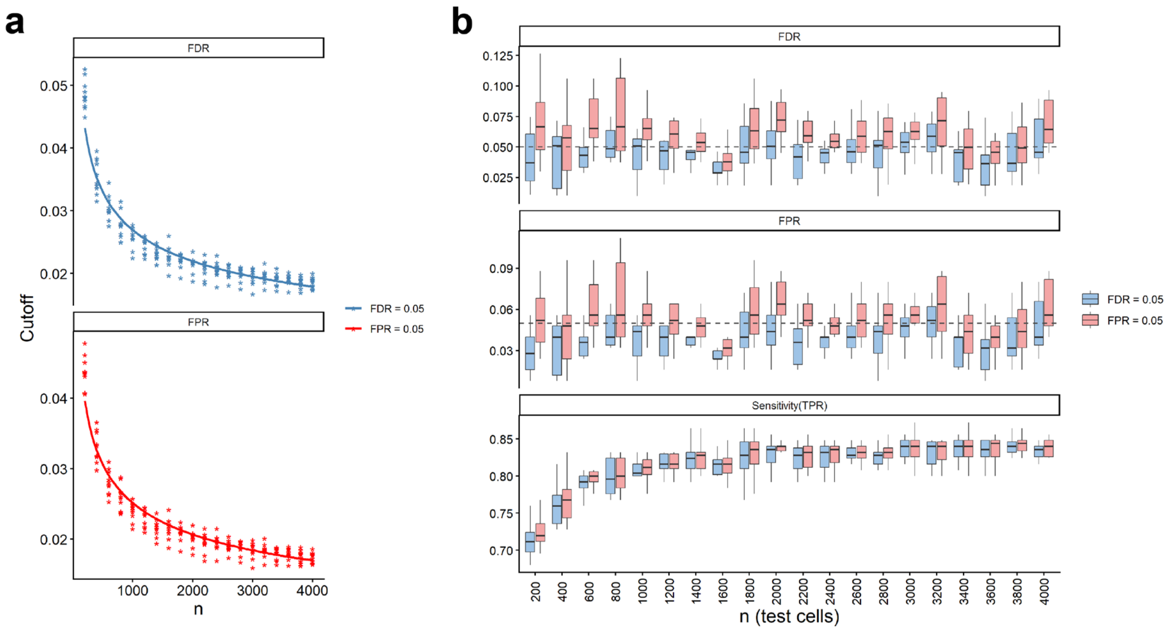

**Supplementary Figure 6.** Learned cutoffs for asymmetric association are stable with increasing sample size.

(**a**) Cutoffs estimated by Algorithm 2 for FDR-controlled (top) and FPR-controlled (bottom) testing at $\alpha = 0.05$ across increasing numbers of cells $n$. Points show individual replicate estimates, and lines show the fitted power-law cutoff function $g(n) = c_1 n^{\{-c_2\}}$. (**b**) Realized FDR, FPR, and sensitivity (TPR) on held-out test cells when the learned cutoffs are applied at each $n$. Boxes show the distribution across replicate datasets for FDR-controlled (blue) and FPR-controlled (pink) cutoffs, and dashed lines mark the 0.05 target.

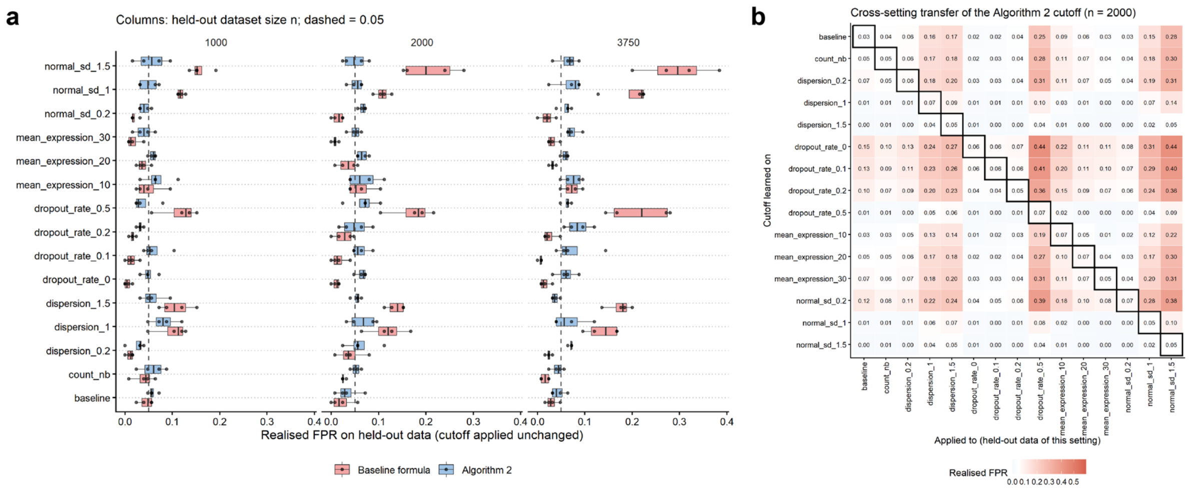

**Supplementary Figure 7.** Cutoffs must be relearned when the simulation parameters change.

(a) Realized FPR on held-out data for cutoffs learned for ChatterjeeKNN under each simulation setting (rows) and applied unchanged to held-out datasets of size $n$ = 1000, 2000, and 3750 (columns). Settings vary the noise standard deviation (normal_sd), mean expression, dropout rate, dispersion, and count distribution (count_nb), relative to the baseline. Cutoffs were estimated either with Algorithm 2 (blue) or with the baseline formula (pink, the fitted FPR-criterion function $g(n) \ = \ c_1 \ n^{\{-c_2\}}$; Supplementary Fig. 6a). Dashed lines mark the 0.05 target. (b) Cross-setting transfer of the Algorithm 2 cutoff at $n$ = 2000. Rows show the setting in which the cutoff was learned, columns show the setting of the held-out data it was applied to, and cell values give the realized FPR. Boxed diagonal cells indicate matched settings.

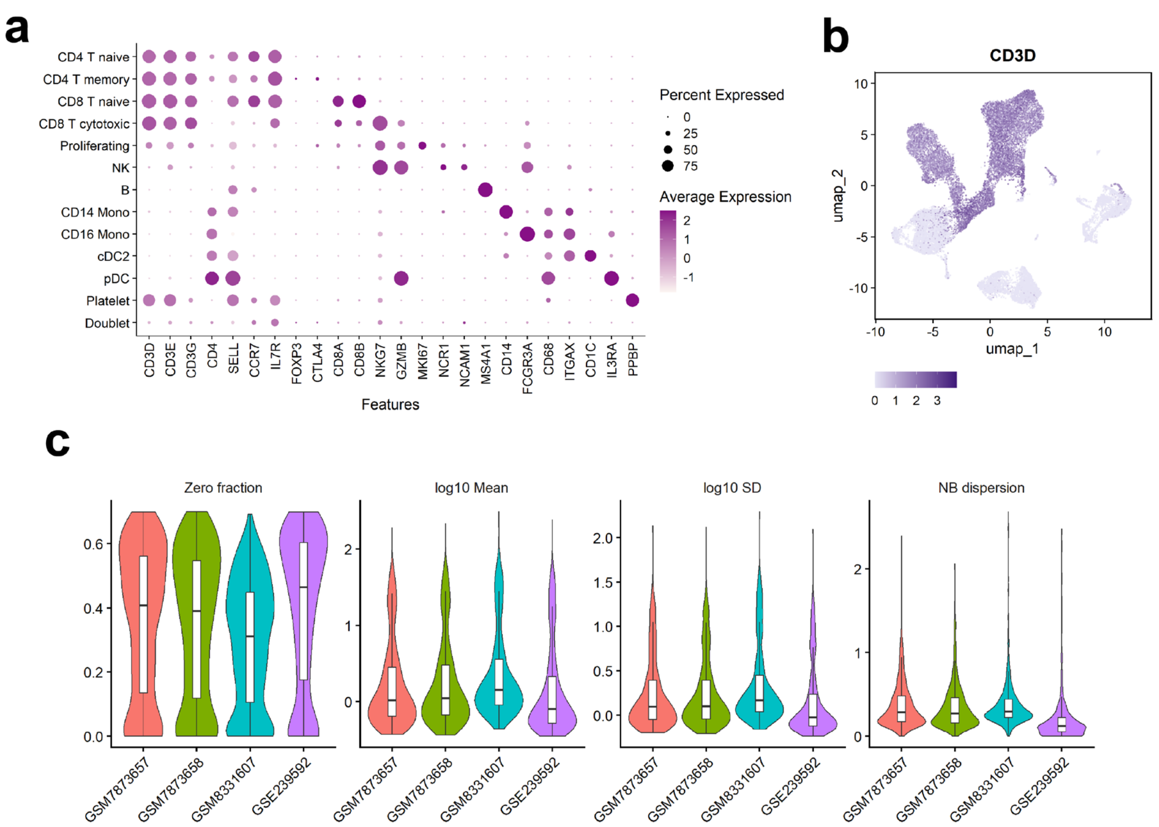

**Supplementary Figure 8.** Cell type annotation and data distributions of the four PBMC datasets.

(**a**) Dot plot of canonical marker genes across the annotated PBMC cell types. Dot size indicates the percentage of cells expressing each gene, and color indicates average expression. (**b**) UMAP of the integrated datasets colored by CD3D expression, the marker used to define T cells. (**c**) Distributions of per-gene zero fraction, log10 mean, log10 standard deviation, and negative binomial (NB) dispersion in T cells for each dataset (GSM7873657, GSM7873658, GSM8331607, and GSE239592). The datasets have similar distributions, supporting transfer of a cutoff learned on one dataset to the others.

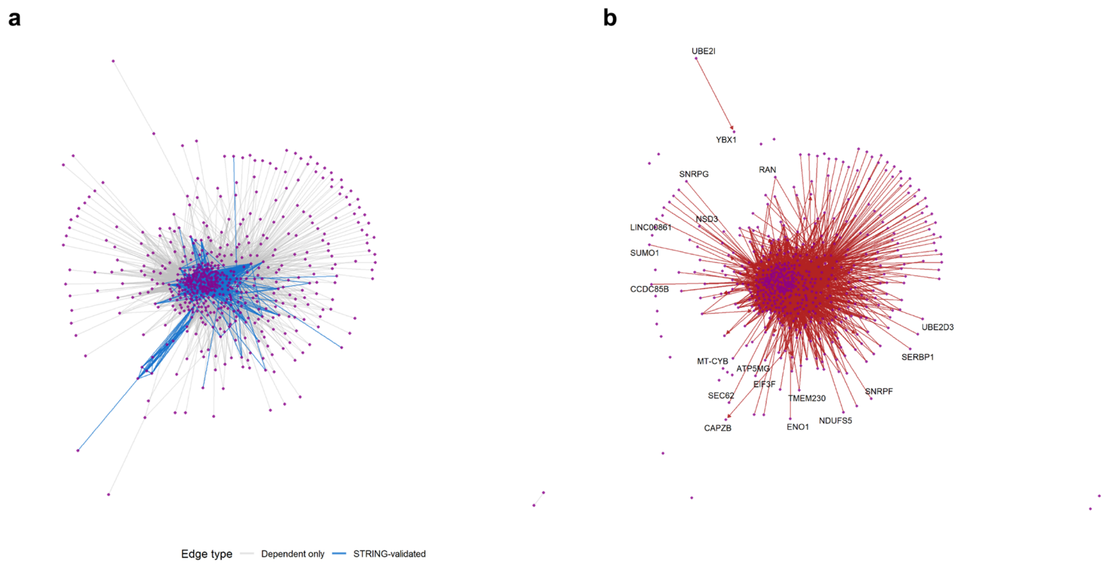

**Supplementary Figure 9.** Dependency and asymmetric association networks inferred from the real PBMC data.

(a) Network of dependent gene pairs identified by the dependency test in the GSM8331607 T cells, using the cutoff learned on GSE239592. Edges also present in the STRING database are shown in blue, and all other dependent pairs are shown in gray. (b) Subnetwork of gene pairs with significant asymmetric association, in which arrows point from the predictor gene to the target gene. Genes at the ends of the directed edges are labeled.

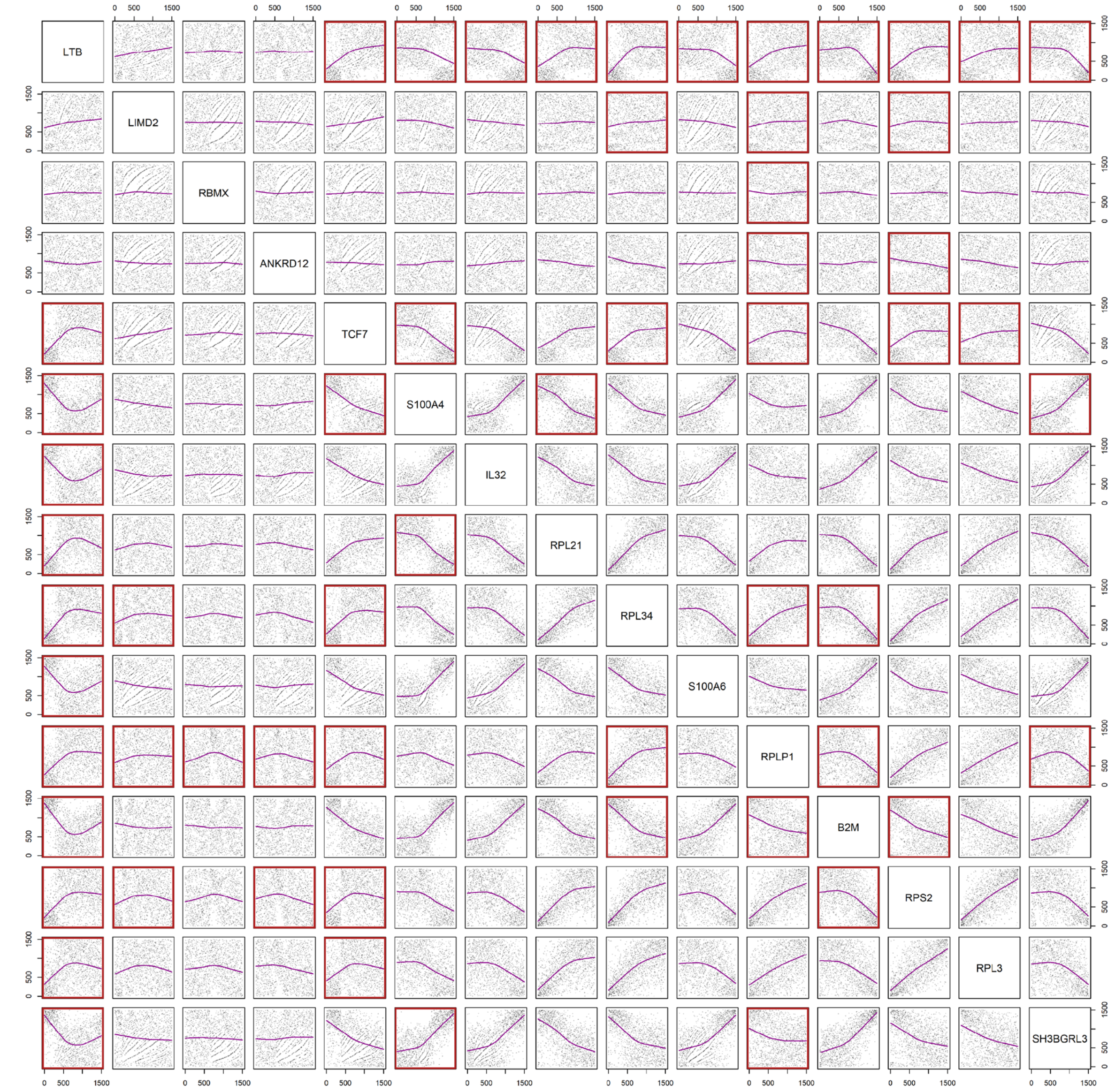

**Supplementary Figure 10.** Pairwise scatter plots of ranked gene expression for genes with significant asymmetric associations in GSM8331607.

Each panel shows the rank-transformed expression of the column gene against the row gene across T cells, with a LOESS fit in purple. Red boxes mark gene pairs with a significant asymmetric association; genes are those in the ten strongest asymmetric pairs.
